# An efficient cognitive–sensorimotor performance state with pupil–heart synchrony independent of subjective flow during esports play

**DOI:** 10.64898/2026.01.15.699797

**Authors:** Shohei Dobashi, Shion Takahashi, Takafumi Yamaguchi, Yukina Tachibana, Ryonosuke Kasahara, Ren Takamizawa, Takashi Matsui

## Abstract

Modern digital activities, such as esports, combine intense cognitive demands with limited whole-body movement, weakening calibration between subjective awareness and functional states. Cognitive deterioration during prolonged esports precedes subjective fatigue, suggesting subjective flow likewise fails to reveal efficient cognitive–sensorimotor performance states. Although efficient cognitive–sensorimotor performance may involve central–autonomic coordination reflected in intrapersonal pupil–heart synchrony, the functional role of this coordination remains largely unknown. We hypothesized that, during esports play, central– autonomic coordination contributes to an efficient cognitive–sensorimotor performance state that subjective flow fails to capture. Fourteen young men completed sessions involving watching a fighting game, playing against a computer-controlled opponent, and competing against another person. We assessed pupil diameter, heart rate, subjective flow, subjective workload, and objective in-game performance. Esports competition increased subjective flow without improving objective in-game performance and reduced combat efficiency, demonstrating a subjectively optimal experience need not signal high in-game performance efficiency. Data-driven clustering identified a low-reactivity state combining superior objective in-game performance with lower subjective workload. This state showed selectively sustained pupil–heart synchrony. Reduced workload partly accounted for the association between pupil– heart synchrony and objective in-game performance. Our findings suggest that individuals fail to recognize periods of high in-game performance efficiency, as cognitive deterioration escapes awareness during prolonged esports. This efficiency is associated with pupil–heart synchrony, supporting the hypothesis that central–autonomic coordination contributes to an efficient cognitive–sensorimotor performance state. Pupil–heart synchrony may therefore serve as a mechanism-based biomarker for detecting efficient states beyond awareness, optimizing objective task performance and the timing of recovery in demanding digital environments

## Introduction

Across human history, from hunting and gathering to contemporary digital work, human activity has relied on cognitive–sensorimotor functions that enable individuals to perceive and predict external events and to select and execute appropriate actions (1, 2). In dynamic whole-body activities, rich bodily feedback helps keep subjective experience coupled to current functional capacity (3, 4). Modern digital activity, by contrast, can disrupt this coupling by allowing agency to be experienced through a virtual body or interface despite limited corresponding whole-body movement (5, 6). Our previous work demonstrated that cognitive decline during prolonged esports preceded subjective fatigue and proposed that enjoyment and potentially flow could combine with low physicality to mask this decline (7). It remains unknown whether the same miscalibration extends to favorable states in which a subjectively optimal experience does not reliably signal high in-game performance efficiency. Both identifying the physiological mechanisms underlying cognitive–sensorimotor performance states that are hidden from subjective awareness and deriving biomarkers from those mechanisms are therefore essential for reconciling optimization of objective task performance with protection from overactivity in modern digital environments (8, 9).

Esports are a representative form of modern cognitive activity in cyberspace, combining intense cognitive–sensorimotor demands with limited whole-body movement (10–12). Esports competition also engages reward-related neural processes that can sustain enjoyment and absorption (13). Within this setting, subjective flow is widely regarded as an optimal experience characterized by deep concentration, absorption, enjoyment, and perceived control (14). Yet flow is not a measure of objective in-game performance. Across tasks, its relationship with objective task performance is inconsistent and depends on cognitive resources, skill, and task experience (15–17); in games, self-reported flow may reflect performance relative to expectations more strongly than the objective in-game performance level itself (17). Flow may remain high even when in-game performance efficiency is low, making esports an ideal setting in which to examine cognitive–sensorimotor performance states that subjective experience alone cannot identify.

Expert esports players exhibit superior cognitive and perceptual–motor abilities that support rapid and accurate in-game performance (18, 19). However, expertise is a trait, whereas in-game performance efficiency varies from moment to moment: even within the same player, acquired abilities can be expressed efficiently or inefficiently across contexts. Their expression depends not only on central processes supporting perception, decision-making, and motor control, but also on autonomic processes that regulate arousal and cardiovascular readiness in response to task demands (2, 20–22). Accordingly, intact coordination among physiological systems within the individual may support a cognitive–sensorimotor performance state in which superior objective in-game performance is achieved with lower subjective workload. We refer to this combination as greater in-game performance efficiency. Disruption of this coordination may instead produce an inefficient state (22–24). Such a state may not be reliably captured by subjective flow (15–17).

One manifestation of such central–autonomic coordination is the temporal coupling between brain and cardiac activity, commonly described as brain–heart coordination. Brain– heart coordination is disrupted in Parkinson’s disease (25), and reductions in this coordination accompany declining subjective focus and objective task performance during sustained attention (24). These findings identify central–autonomic coordination as a candidate mechanism linking arousal regulation to task performance and intrapersonal physiological synchrony as an observable manifestation of this coordination. Direct assessment of brain–heart coordination using electroencephalography, functional near-infrared spectroscopy, or functional magnetic resonance imaging, however, requires specialized equipment that constrains naturalistic game and esports play and continuous real-world monitoring (26). A practical biomarker requires signals that capture central and autonomic dynamics without interrupting ongoing task execution.

Under constant luminance, pupil diameter is a noninvasive index of ascending arousal-system activity (27, 28). In our previous study, pupil constriction tracked cognitive decline during prolonged esports even when subjective fatigue did not (7), demonstrating that pupil dynamics can reveal functional deterioration that escapes awareness. Heart rate provides a complementary index of peripheral autonomic and cardiovascular readiness (29), and both signals can be monitored continuously using wearable devices (30). Moreover, pupil-linked and heartbeat-linked arousal make complementary contributions to behavioral performance, and their combined assessment predicts behavior better than either signal alone (31). The temporal coordination of pupil diameter and heart rate, here termed pupil–heart synchrony, may reflect central–autonomic coordination between pupil-linked central arousal and heart-rate-linked autonomic cardiovascular readiness during demanding digital activity. This coordination could contribute to an efficient cognitive–sensorimotor performance state, while pupil–heart synchrony provides a physiological basis for identifying that state beyond subjective flow. Pupil–heart synchrony may therefore serve as a scalable, mechanism-based biomarker for distinguishing efficient from inefficient cognitive–sensorimotor performance states.

Accordingly, we hypothesized that, during game and esports play, central–autonomic coordination reflected in intrapersonal pupil–heart synchrony contributes to an efficient cognitive–sensorimotor performance state that subjective flow fails to capture. To test this hypothesis across contexts that vary in opponent type and interactivity (32, 33), we assessed pupil diameter, heart rate, subjective flow, subjective workload, and objective in-game performance across three conditions: watching a fighting game, playing against a computer-controlled opponent, and competing against another person. Within this design, we made three predictions. First, competitive play would elevate subjective flow without a commensurate improvement in objective in-game performance. Second, pupil–heart synchrony would distinguish efficient from inefficient cognitive–sensorimotor performance states beyond subjective flow. Third, the state exhibiting pupil–heart synchrony would combine superior objective in-game performance with lower subjective workload, indicating greater in-game performance efficiency. As part of this prediction, reduced workload would statistically mediate the association between pupil–heart synchrony and objective in-game performance.

## Results

### Competitive play increases subjective flow without improving objective in-game performance

To address the study aims, fourteen healthy young adult men completed repeated sessions of a one-versus-one fighting game (Super Smash Bros. Ultimate, Nintendo, Japan) under three conditions: passively watching the game (Watch), playing against a computer-controlled opponent (Game), and competing against a human opponent (Esports) (Figure 1A). The Watch condition served as a passive baseline with comparable game-related visual input without game play. This design supported two complementary levels of analysis: condition-level comparisons to characterize context-dependent effects, and session-level analyses that first used unsupervised k-means clustering of joint pupil–heart reactivity to identify physiological states and then applied cross-recurrence quantification analysis (CRQA) to quantify nonlinear temporal coordination between the pupil and heart-rate time series beyond their mean reactivity (Figure 1B).

**Figure 1.**
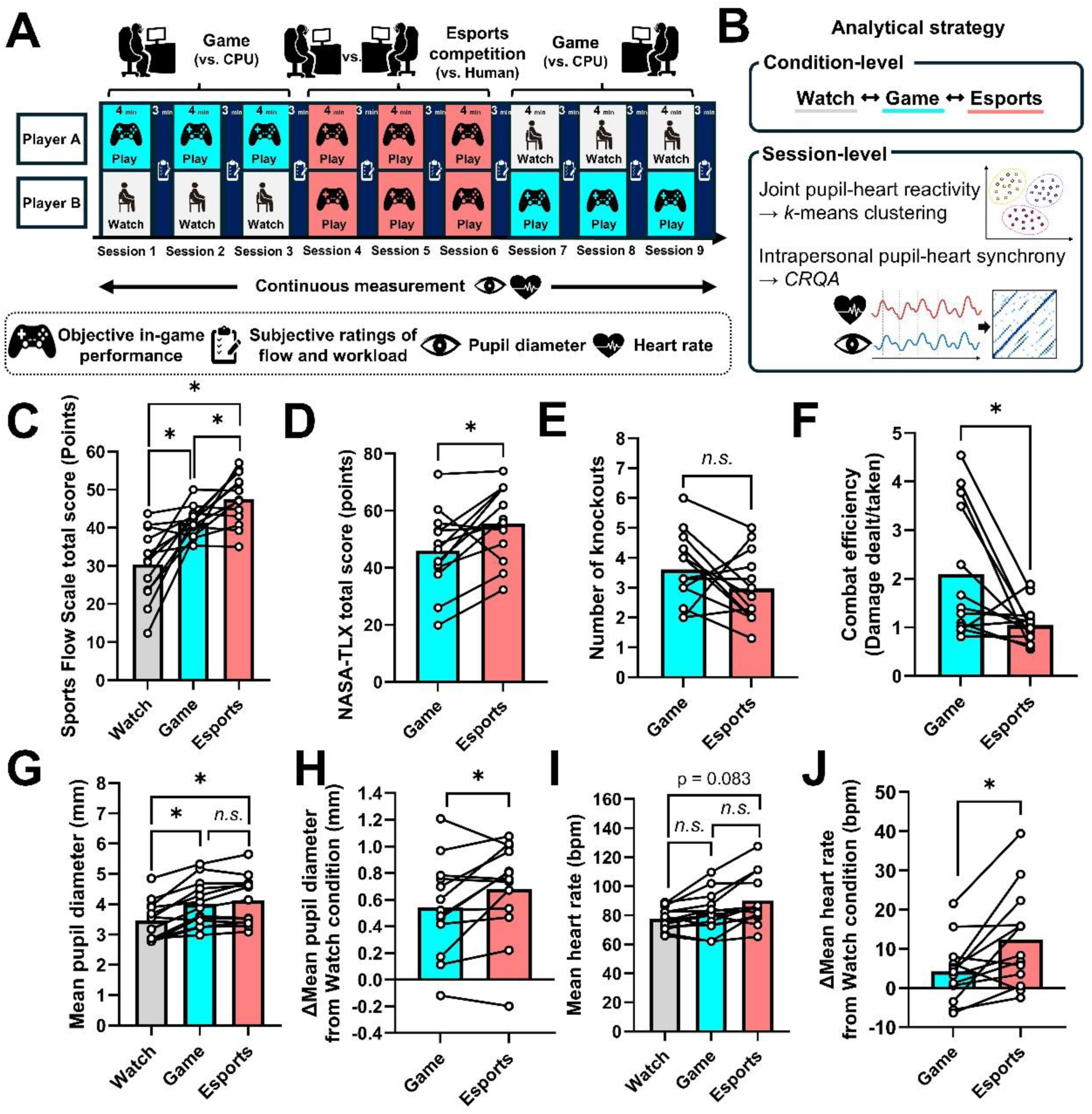
Esports competition increases subjective flow without improving objective in-game performance. **A.** Overview of the experimental protocol. Across nine sessions, paired participants completed three conditions: passively watching the game (Watch), playing against a computer-controlled opponent (Game), and competing against each other (Esports). **B.** Analytical framework of this study. This design supported two complementary levels of analysis: condition-level comparisons to characterize context-dependent effects, and session-level analyses that first used unsupervised k-means clustering of joint pupil–heart reactivity to identify physiological states and then applied cross-recurrence quantification analysis (CRQA) to quantify nonlinear temporal coordination between the pupil and heart-rate time series beyond their mean reactivity. **C.** Sports Flow Scale (SFS) total score. **D.** NASA Task Load Index (NASA-TLX) total score. **E–F.** Objective in-game performance outcomes: number of knockouts (**E**) and combat efficiency, calculated as damage dealt divided by damage taken (**F**). **G.** Mean pupil diameter. **H.** Change in mean pupil diameter relative to the Watch condition. **I.** Mean heart rate. **J.** Change in mean heart rate relative to the Watch condition. For each participant, values were averaged across the three sessions within each condition. Bars represent group means, points represent individual participants, and lines connect within-participant values across conditions. Condition effects in **C, G,** and **I** were evaluated using one-way repeated-measures ANOVA followed by Bonferroni-corrected paired comparisons. Comparisons in **D–F, H,** and **J** were performed using paired *t* tests. *p* < 0.05; n.s., not significant. Detailed statistical results are provided in Supplementary Tables 2 and 3.

We first examined whether competitive play increased subjective flow without a commensurate improvement in objective in-game performance. Subjective flow was assessed using the Sports Flow Scale (SFS). SFS total scores differed across conditions (p < 0.05, η²p = 0.637), with higher scores during active play and the highest scores during human competition (Esports; Cohen’s dz = 0.918 vs. Game) (Figure 1C). Multiple dimensions of flow were also modulated by active play, although the changes were not uniform across subscales (Supplementary Figure 1A; Supplementary Tables 2 and 3). Subjective workload was assessed using the NASA Task Load Index (NASA-TLX; see Appendix). Compared with game play, human competition significantly increased NASA-TLX total scores (p < 0.05, Cohen’s dz = 0.796) (Figure 1D), along with selected workload components (Supplementary Figure 1B; Supplementary Tables 2 and 3). Although group-level SFS and NASA-TLX scores were highest during the Esports condition, both measures varied substantially across participants, with some participants reporting higher scores during Game.

Objective in-game performance was assessed using the number of knockouts, combat efficiency, damage dealt, damage taken, and attacking accuracy; comparisons of these metrics between winning and losing sessions are shown in Supplementary Figure 2A–E. Combat efficiency, an objective in-game performance metric integrating offensive output and defensive cost, was calculated as the ratio of damage dealt to damage taken. Compared with Game, Esports reduced combat efficiency (p = 0.017, dz = 0.768), whereas the number of knockouts did not differ significantly between conditions (p = 0.155, dz = 0.421) (Figure 1E and F). The Esports condition also reduced damage dealt and increased damage taken (damage dealt: p = 0.037, dz = 0.652; damage taken: p = 0.016, dz = 0.780), whereas attacking accuracy did not differ between conditions (p = 0.648, dz = 0.133) (Supplementary Figure 3A–C). Thus, human competition reduced combat efficiency by decreasing offensive output and increasing defensive cost without increasing knockouts or attacking accuracy. Consistent with this dissociation, SFS total scores were not associated with either knockouts or combat efficiency after accounting for repeated observations within participants (Supplementary Figure 4A and B).

Physiological responses also varied across play conditions. Pupil diameter increased during active play compared with the Watch condition (p < 0.0001, η²p = 0.729), and heart rate showed a significant overall condition effect (p < 0.001, η²p = 0.457) (Figure 1G and I). Analyses relative to the watching baseline confirmed that both pupil diameter and heart rate were modulated during active play (pupil diameter: p = 0.037, dz = 0.651; heart rate: p = 0.009, dz = 0.871) (Figure 1H and J). Both physiological responses exhibited substantial interindividual variability.

Together, competitive play increased subjective flow and subjective workload but did not improve objective in-game performance and instead reduced combat efficiency. Pupil-diameter and heart-rate responses also varied substantially across individuals, motivating the examination of joint physiological response profiles.

### Joint pupil–heart reactivity identifies an efficient cognitive–sensorimotor performance state beyond subjective flow

Because subjective flow did not reliably track objective in-game performance, we next tested whether joint pupil–heart reactivity identified distinct physiological profiles associated with cognitive–sensorimotor performance states. Changes in pupil diameter and heart rate during active play were moderately correlated but not fully aligned (Pearson’s r = 0.396, p < 0.001). We classified active play sessions according to their joint reactivity using two-dimensional k- means clustering of changes from the watching baseline (Figure 2A).

**Figure 2.**
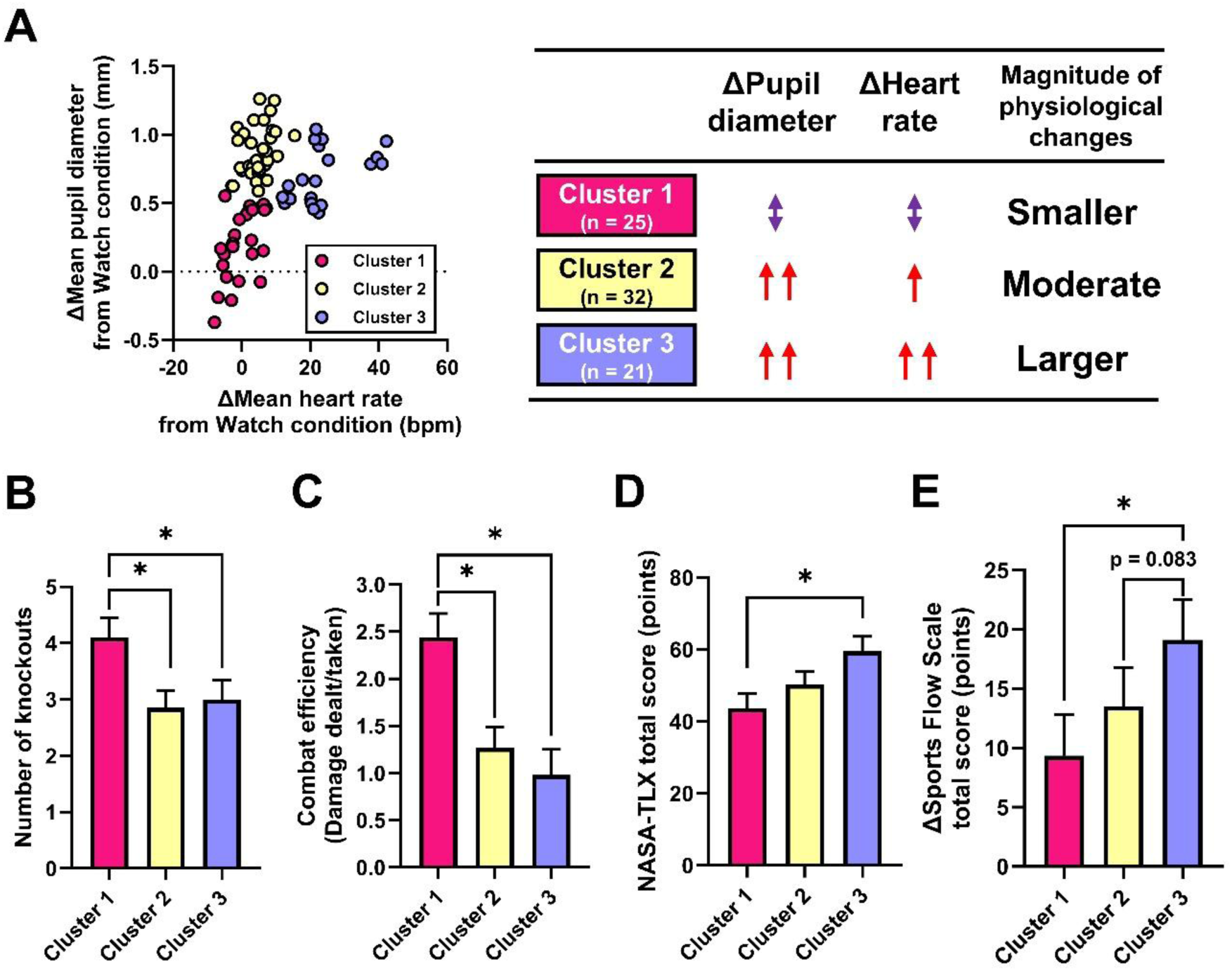
Joint pupil–heart reactivity identifies an efficient cognitive–sensorimotor performance state not captured by subjective flow. **A.** Two-dimensional *k*-means clustering (*k* = 3) of changes in mean pupil diameter and heart rate during active play relative to each participant’s Watch baseline. Each point represents one active play session. The resulting clusters were designated as low-, intermediate-, and high-reactivity physiological profiles. **B–C.** Objective in-game performance outcomes across clusters: number of knockouts (**B**) and combat efficiency, calculated as damage dealt divided by damage taken (**C**). **D.** Subjective workload assessed using the NASA Task Load Index (NASA-TLX) total score. **E.** Change in subjective flow assessed using the Sports Flow Scale (SFS) total score relative to the Watch condition. Bars in **B–E** represent estimated marginal means ± SEM derived from linear mixed-effects models with participant identity as a random intercept. Significant cluster effects were followed by Bonferroni-corrected contrasts of estimated marginal means. *p* < 0.05; n.s., not significant. Detailed statistical results are provided in Supplementary Tables 4, 6, and 8.

The 78 active play sessions included in the clustering yielded three physiological response profiles. Cluster 1 (n = 25) exhibited minimal or reduced changes in both pupil diameter and heart rate, Cluster 2 (n = 32) exhibited moderate increases, and Cluster 3 (n = 21) exhibited relatively large increases in both measures (Figure 2A). After accounting for repeated observations within participants, play condition was not significantly associated with physiological cluster membership (multinomial logistic regression with standard errors clustered by participant, Wald χ²(2) = 4.09, p = 0.130; Supplementary Figure 5A). Similarly, win–loss outcomes did not differ significantly across clusters (mixed-effects logistic regression, Wald χ²(2) = 3.44, p = 0.179; Supplementary Figure 5B). Together, these results suggest that the physiological clusters were unlikely to have been driven solely by play condition or win–loss outcome.

We next examined whether objective in-game performance and subjective workload differed across clusters. The number of knockouts and combat efficiency were significantly higher in Cluster 1 than in Cluster 3 (knockouts: p = 0.044, b = 1.11; combat efficiency: p < 0.001, b = 1.46) (Figure 2B and C). Cluster 1 also showed more favorable damage dealt, damage taken, and attacking accuracy than Cluster 3 (Supplementary Figure 6A–C).

Conversely, subjective workload (NASA-TLX total score) was significantly higher in Cluster 3 than in Cluster 1 (p = 0.005, b = 15.79) (Figure 2D), with greater mental demand in Cluster 3 (Supplementary Figure 6D). These cluster differences in objective in-game performance were generally preserved after adjustment for play condition (Supplementary Table 5). Cluster 1 combined superior objective in-game performance with lower subjective workload, indicating greater in-game performance efficiency and operationally identifying it as an efficient cognitive–sensorimotor performance state.

Subjective flow did not identify this efficient state. Absolute SFS total scores did not differ significantly across clusters (Supplementary Figure 7A). Changes in SFS total scores nevertheless differed across clusters, with a greater increase in Cluster 3 than in Cluster 1 (p = 0.004, b = 9.77) (Figure 2E; Supplementary Figure 7C). Flow-subscale profiles were also heterogeneous: “focus on current task” was higher in Cluster 3, whereas “loss of self-consciousness” was lower in Cluster 1 (Supplementary Figure 7B and D). Thus, the largest increase in subjective flow occurred in the high-reactivity state rather than in the state with the greatest in-game performance efficiency.

Together, joint pupil–heart reactivity identified an efficient cognitive–sensorimotor performance state characterized by superior objective in-game performance and lower subjective workload. This state was not characterized by greater subjective flow and could not be identified from subjective flow alone.

### The efficient cognitive–sensorimotor performance state exhibits selectively sustained pupil–heart synchrony

We next examined whether the efficient cognitive–sensorimotor performance state exhibited distinct temporal coordination between pupil and heart dynamics. Neither maximum cross-correlation nor its corresponding time lag differed across clusters (Supplementary Figure 8A and B), indicating that conventional linear coupling did not distinguish the identified states. We used cross-recurrence quantification analysis (CRQA) to characterize more complex temporal coordination between pupil diameter and heart rate.

Under a fixed recurrence rate, Cluster 1 exhibited a significantly smaller recurrence radius and a longer maximum diagonal line length than Cluster 3 (recurrence radius: p = 0.020, b = −2.72; maximum diagonal line length: p = 0.004, b = 1.74) (Figure 3B and E). These features indicate more tightly constrained pupil–heart dynamics and longer episodes of sustained temporal coordination. Consistent with the smaller recurrence radius, Cluster 1 exhibited a more compact and constrained distribution in pupil–heart state space (Supplementary Figure 9A).

**Figure 3.**
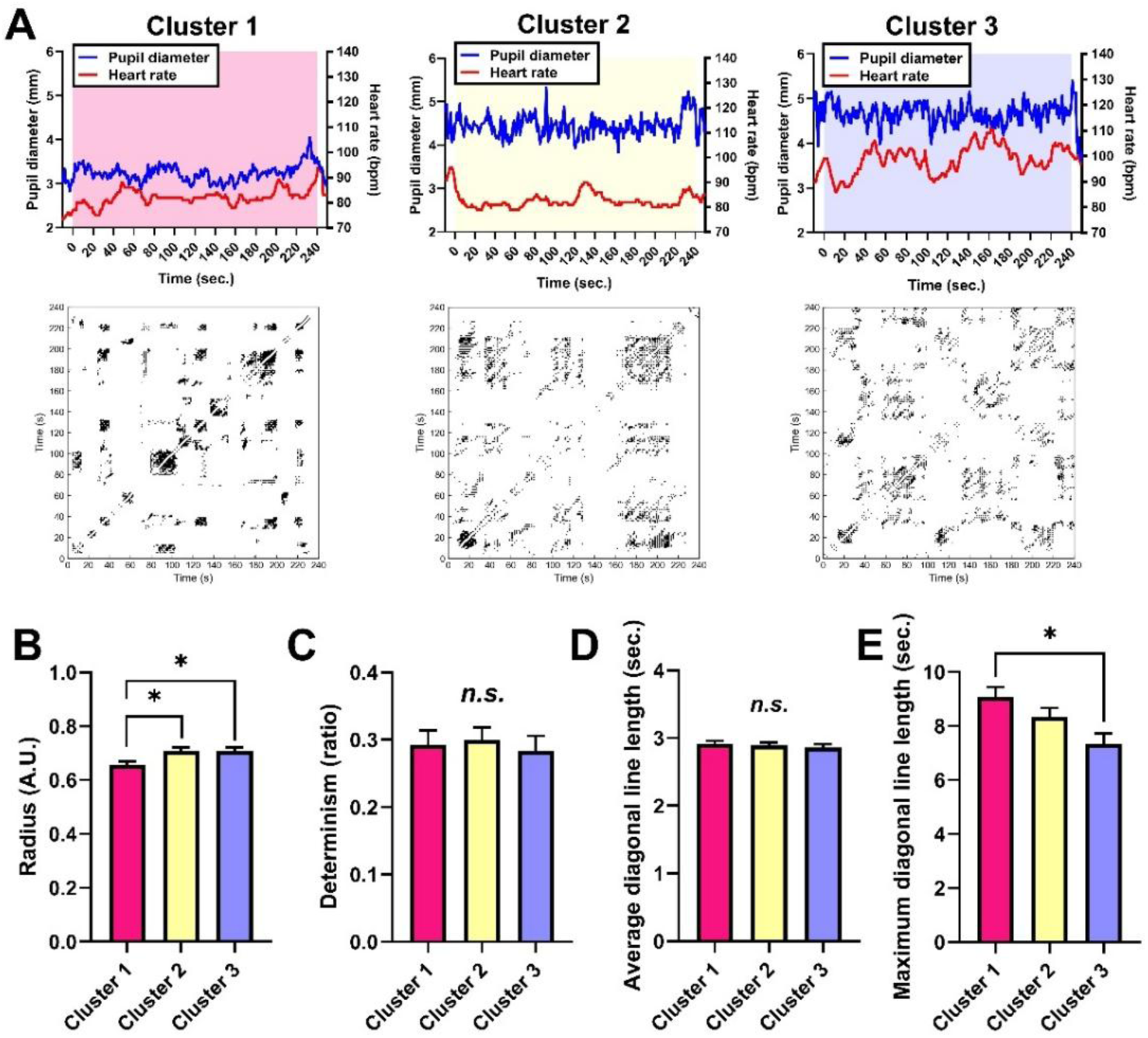
The efficient cognitive–sensorimotor performance state exhibits selectively sustained pupil–heart synchrony. **A.** Representative pupil-diameter and heart-rate time series and the corresponding cross-recurrence plots from each physiological cluster. Diagonal structures in the recurrence plots indicate episodes of similar temporal evolution between pupil and heart-rate dynamics, with longer diagonal structures indicating more sustained temporal coordination. **B–E.** Cross-recurrence quantification analysis (CRQA) indices calculated under a fixed recurrence rate of 5%: recurrence radius (**B**), determinism (**C**), average diagonal line length (**D**), and maximum diagonal line length (**E**). A smaller recurrence radius indicates more tightly constrained joint pupil–heart dynamics, whereas a longer maximum diagonal line indicates a longer episode of sustained temporal coordination. Bars represent estimated marginal means ± SEM derived from linear mixed-effects models with participant identity as a random intercept. Significant cluster effects were followed by Bonferroni-corrected contrasts of estimated marginal means. \**p* < 0.05; n.s., not significant. Additional CRQA results are provided in Supplementary Figure 9 and Supplementary Table 10.

Other CRQA indices did not differ significantly across clusters. These included determinism and average diagonal line length (Figure 3C and D; Supplementary Table 10), as well as diagonal entropy, laminarity, average and maximum vertical line lengths, and vertical entropy (Supplementary Figure 9B–F; Supplementary Table 10).

Thus, the efficient cognitive–sensorimotor performance state was characterized not by uniformly stronger conventional linear coupling but by more tightly constrained pupil–heart dynamics and selectively sustained temporal synchrony.

### Reduced subjective workload statistically mediates the association between pupil–heart synchrony and objective in-game performance

Finally, we tested whether subjective workload statistically mediated the association between pupil–heart synchrony and objective in-game performance. Pupil–heart coordination was indexed by CRQA recurrence radius, with a smaller radius indicating more tightly constrained joint pupil–heart dynamics under the fixed recurrence rate. Mediation models specified recurrence radius as the predictor, NASA-TLX total score as the mediator, and objective in-game performance metrics as the outcomes (Figure 4, Supplementary Figure 10).

**Figure 4.**
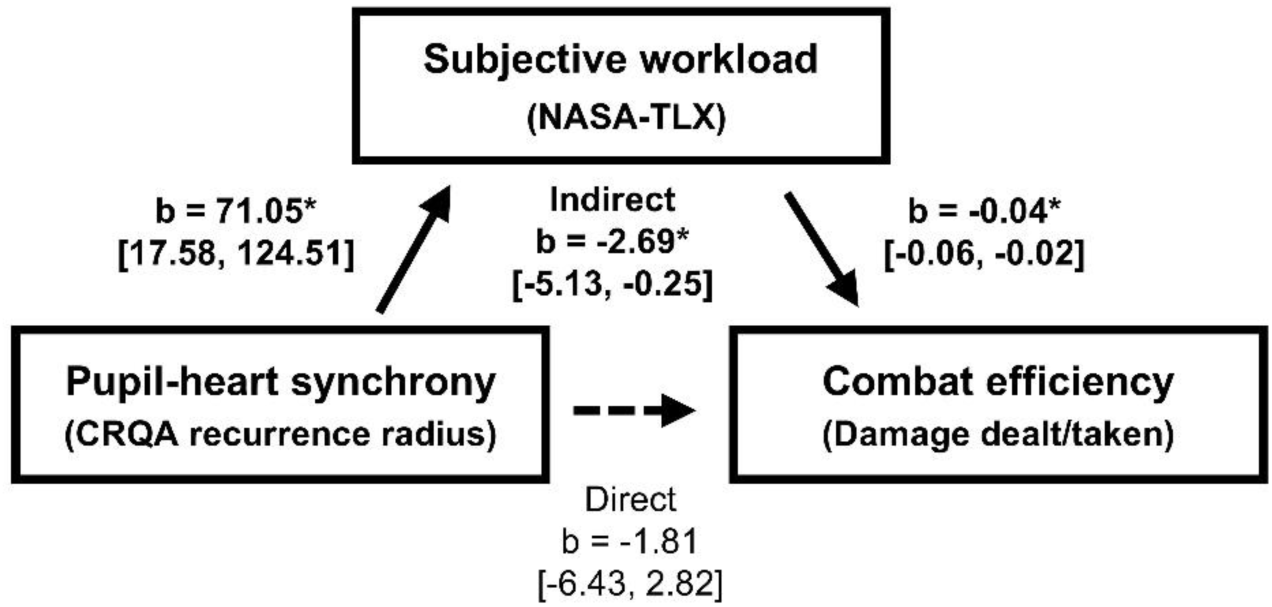
Reduced subjective workload statistically mediates the association between pupil–heart synchrony and objective in-game performance. Mediation models testing whether subjective workload statistically accounted for part of the association between pupil–heart coordination and objective in-game performance. CRQA recurrence radius was entered as the predictor, NASA Task Load Index (NASA-TLX) total score as the mediator, and combat efficiency as the outcome. A smaller recurrence radius indicates more tightly constrained joint pupil–heart dynamics under the fixed recurrence-rate criterion. Indirect effects were calculated as the product of the predictor-to-mediator and mediator-to-outcome coefficients. Values are presented as unstandardized coefficients (*b*) with 95% confidence intervals. *p* < 0.05. Detailed statistical results are provided in Supplementary Table 11.

A smaller recurrence radius was associated with lower subjective workload, which in turn was associated with greater combat efficiency. This pathway produced a significant indirect effect on combat efficiency (indirect effect b = −2.69, 95% CI [−5.13, −0.25]) (Figure 4), indicating that reduced subjective workload statistically accounted for part of the association between more tightly constrained pupil–heart dynamics and greater in-game performance efficiency.

Significant indirect effects through subjective workload were also observed for the number of knockouts (b = −2.59, 95% CI [−5.00, −0.17]), damage dealt (b = −324.12, 95% CI [−612.48, −35.76]), and damage taken (b = 193.47, 95% CI [24.19, 362.75]), whereas the indirect effect for attacking accuracy was not significant (b = −9.27, 95% CI [−20.41, 1.86]) (Supplementary Figure 10A–D). Among these outcomes, only damage taken retained a significant direct association with recurrence radius after accounting for subjective workload (direct effect b = 556.65, 95% CI [65.19, 1048.10]).

Together, reduced subjective workload statistically accounted for part of the association between pupil–heart synchrony and multiple objective in-game performance outcomes. The additional direct association with damage taken indicates that pupil–heart synchrony may also relate to the defensive component of objective in-game performance independently of subjective workload.

## Discussion

The present findings support the hypothesis that, during game and esports play, central– autonomic coordination reflected in intrapersonal pupil–heart synchrony contributes to an efficient cognitive–sensorimotor performance state that subjective flow fails to capture. Esports competition increased subjective flow without improving objective in-game performance and reduced combat efficiency (Figure 1C, E, and F), showing that a subjectively optimal experience does not necessarily signal high in-game performance efficiency. Data-driven clustering of pupil-diameter and heart-rate responses instead identified a low-reactivity state with superior objective in-game performance and lower subjective workload (Figure 2A–D), indicating greater in-game performance efficiency. This state also showed selectively sustained pupil–heart synchrony, and reduced workload statistically accounted for part of the association between pupil–heart synchrony and objective in-game performance (Figure 3B and E; Figure 4). Just as cognitive deterioration during prolonged esports can escape subjective awareness (7), the present study shows that individuals may also fail to recognize periods of particularly high in-game performance efficiency, revealing the two-sided nature of subjective–objective miscalibration in modern digital activity.

### Subjective flow dissociates from objective in-game performance

The dissociation between subjective flow and objective in-game performance was evident across condition, session, and cluster analyses. Human competition increased subjective flow without improving objective in-game performance and reduced combat efficiency (Figure 1C, E, and F). This pattern is consistent with evidence that playing against a human-controlled opponent elicits greater subjective flow than playing against a computer-controlled opponent (33). Session-level flow scores were also unrelated to knockouts or combat efficiency (Supplementary Figure 4A and B). Across physiological clusters, objective in-game performance differed, whereas absolute subjective flow scores did not (Figure 2B and C; Supplementary Figures 6A–C and 7A). Cluster 3 showed the largest increase in flow without a corresponding advantage in objective in-game performance (Figure 2B, C, and E). These physiological profiles were not explained by play condition or win–loss outcome (Supplementary Figure 5). Together, these analyses indicate that subjective flow tracked the competitive context more closely than objective in-game performance.

One explanation for this dissociation concerns what subjective flow represents. Flow is characterized by absorption in the ongoing experience and effortless attention (14, 15), while its association with objective task performance is small to moderate across studies (16). Flow has also been examined as a distinctive experience during esports play and as a coordinated neurophysiological state during team performance (34, 35). In game tasks, self-reported flow may reflect better-than-expected performance more strongly than actual objective task performance (17). Subjective flow may instead represent how favorably ongoing performance is appraised relative to expectations rather than the efficiency of the underlying cognitive– sensorimotor performance state.

The convergence of these condition-level, session-level, and cluster-level dissociations lends indirect support to an embodied-calibration account. Although bodily feedback was not manipulated, participants met intense cognitive–sensorimotor demands through a digital interface with limited whole-body movement. Such configurations can preserve agency and immersion despite limited corresponding movement (5, 6), while providing less bodily feedback with which to calibrate subjective experience against the current functional state (3, 4). This pattern is consistent with the possibility that strong flow can coexist with inaccurate awareness of the efficiency of the underlying cognitive–sensorimotor performance state.

The unrecognized efficiency observed here represents a favorable counterpart to the problem identified in our previous study. Whereas cognitive decline during prolonged esports preceded subjective fatigue (7), the particularly efficient cognitive–sensorimotor performance state observed here was not captured by subjective flow. Subjective–objective miscalibration in modern digital activity is therefore two-sided, encompassing failures to recognize either deterioration in functional state or particularly high in-game performance efficiency. The implications of these failures are complementary, as unrecognized deterioration may prolong activity and delay recovery, whereas unrecognized efficiency may hinder identification and reproduction of the physiological conditions that support greater in-game performance efficiency. This calibration problem creates a need for objective physiological markers capable of detecting both unfavorable and favorable cognitive–sensorimotor performance states beyond subjective awareness.

### Pupil–heart synchrony characterizes an efficient cognitive–sensorimotor performance state

Joint pupil–heart dynamics distinguished an efficient cognitive–sensorimotor performance state that subjective flow did not. Data-driven clustering of pupil and heart-rate reactivity identified Cluster 1 as a low-reactivity profile characterized by relatively small changes in pupil diameter and heart rate (Figure 2A). This profile nevertheless combined superior objective in-game performance with lower subjective workload (Figure 2B–D). The convergence of these outcomes operationally defines Cluster 1 as an efficient cognitive–sensorimotor performance state.

CRQA further revealed that this efficient state was characterized by distinct temporal coordination between pupil and heart dynamics. Conventional cross-correlation strength and temporal lag did not differ across clusters (Supplementary Figure 8A and B). By contrast, Cluster 1 exhibited a smaller recurrence radius and a longer maximum diagonal line length than Cluster 3 (Figure 3B and E). Under the fixed recurrence-rate analysis, these features indicate more tightly constrained joint dynamics and longer episodes of sustained temporal coordination. Global recurrence measures, including determinism, average diagonal line length, and laminarity, also did not differ across clusters, as shown in Figure 3C and D and Supplementary Figure 9B–F. Thus, the efficient cognitive–sensorimotor performance state was characterized by selectively sustained and tightly constrained pupil–heart coordination rather than uniformly stronger synchrony. Together with the modest physiological reactivity of Cluster 1, this pattern suggests that in-game performance efficiency may depend more on intrapersonal coordination between central and autonomic responses than on the magnitude of either response alone. This interpretation is consistent with the neural efficiency hypothesis, which proposes that superior performance reflects the efficient allocation of task-relevant resources rather than greater overall resource mobilization (36, 37).

Mediation analysis indicated that lower subjective workload statistically accounted for part of the association between a smaller CRQA radius and objective in-game performance outcomes (Figure 4; Supplementary Figure 10A–C). Damage taken also retained an association with pupil–heart coordination after subjective workload was included in the model. This distinction is consistent with partly different pathways linking physiological coordination to components of objective in-game performance. Damage taken may more directly reflect reactive perceptual–motor processing, including detecting and avoiding an opponent’s attacks (38, 39), whereas knockouts and damage dealt depend more strongly on accumulated judgments, action selection, and offensive decision-making (12, 39). This distinction points to a more direct relationship between pupil–heart coordination and defensive sensorimotor control, while reduced subjective workload statistically accounts for part of its association with broader objective in-game performance.

Together, these findings indicate that pupil–heart synchrony reflects central– autonomic coordination in an efficient cognitive–sensorimotor performance state. Pupil–heart synchrony may therefore serve as a mechanism-based biomarker of that state. The informative physiological pattern was not generalized physiological activation or uniformly stronger coupling, but modest reactivity and sustained central–autonomic coordination associated with superior objective in-game performance and lower subjective workload. This profile is consistent with a physiologically characterized “zone-like” state in which high-level objective in-game performance is expressed with relatively low subjective effort (40).

### Why was pupil–heart synchrony preserved in the efficient state?

The selective CRQA pattern, together with the small task-evoked changes in pupil diameter and heart rate in Cluster 1, constrains the physiological interpretation of the efficient state (Figure 2A, Figure 3B and E, and Supplementary Figure 9A). Cross-correlation strength and temporal lag did not differ across clusters, nor did the other CRQA indices (Supplementary Figure 8A and B, Figure 3C and D, and Supplementary Figure 9B–F). Thus, the efficient cognitive– sensorimotor performance state was not characterized by generalized physiological activation, uniformly stronger synchrony, or a simple shift in temporal lag. Instead, it combined modest physiological reactivity with selectively sustained and tightly constrained pupil–heart coordination.

This pattern is consistent with repeated temporal alignment between pupil-linked central arousal and heart-rate-linked autonomic cardiovascular readiness during behaviorally relevant in-game events. Such coordination could emerge transiently when perceptual, decisional, or motor demands require corresponding central and bodily preparation rather than remaining continuously elevated throughout active play. The diagonal structures in the representative cross-recurrence plots are consistent with this episodic account (Figure 3A). Event-dependent alignment could account for the selective CRQA pattern without requiring stronger overall correlation.

The locus coeruleus–norepinephrine system provides a plausible mechanism for this pattern. Adaptive Gain Theory proposes that moderate tonic locus coeruleus activity supports efficient phasic responses to behaviorally relevant stimuli, whereas high tonic activity attenuates such responses (41). Consistent with this account, smaller baseline pupil diameter has been associated with larger phasic pupil responses and faster reaction times during an oddball task (42). Changes in pupil dilation and P300 amplitude have also implicated the locus coeruleus– norepinephrine system in psychological flow (43). The modest physiological reactivity of Cluster 1 could reflect relatively stable tonic arousal that permitted selective phasic responses to relevant in-game events. Successful expression of these responses may also require the autonomic and cardiovascular systems to enter a corresponding preparatory state (31, 42). Pupil–heart synchrony may consequently reflect repeated alignment of central arousal with bodily readiness rather than continuous coupling throughout active play.

Experimentally induced emotional arousal has also been associated with measure-dependent alterations in brain–heart interactions during immersive virtual reality (44). The greater physiological reactivity of Clusters 2 and 3 may have destabilized central–autonomic coordination, whereas the constrained reactivity of Cluster 1 may have allowed this coordination to remain temporally organized.

Taken together, pupil–heart synchrony characterizes a physiological state in which overall physiological reactivity is constrained while pupil-linked central arousal remains selectively coordinated with autonomic cardiovascular readiness. At a conceptual level, this organization resembles *mushin* and *heijoshin*, which martial arts scholarship describes as freedom from mental fixation together with a calm and undisturbed readiness to respond (45, 46). Accordingly, pupil–heart synchrony offers a testable physiological framework for investigating how constrained arousal and selectively sustained central–autonomic coordination may support these states.

#### Toward mechanism-based monitoring of cognitive–sensorimotor performance states

The present study provides a physiological basis for monitoring cognitive–sensorimotor performance states beyond subjective awareness. A low-reactivity state combined superior objective in-game performance with lower subjective workload (Figure 2A–D) and exhibited selectively sustained pupil–heart synchrony (Figure 3B and E). Reduced workload also statistically accounted for part of the association between pupil–heart synchrony and objective in-game performance (Figure 4; Supplementary Figure 10). Together, the low-reactivity profile, sustained synchrony, and mediation results support the hypothesis that central–autonomic coordination contributes to an efficient cognitive–sensorimotor performance state that subjective flow fails to capture. On this basis, pupil–heart synchrony may serve as a mechanism-based biomarker of that state. Whereas our previous study showed that pupil constriction can detect cognitive deterioration that precedes subjective fatigue during prolonged esports (7), pupil–heart synchrony may detect otherwise unrecognized efficiency. Multimodal monitoring could yield complementary indicators of unfavorable and favorable states that escape subjective awareness.

Such monitoring is technically plausible. Heart rate can be recorded continuously using wearable devices, and real-time pupillometry using camera-based systems has been demonstrated under controlled conditions (47). Biosignal sharing has also been incorporated into online games to augment social presence (48), indicating that physiological feedback can be integrated into digital play. Importantly, the proposed biomarker depends not simply on the magnitude or direction of either physiological response but on their temporal coordination. This feature may distinguish efficient central–autonomic organization from nonspecific physiological arousal. Real-world use will require validation across devices, lighting conditions, head movements, and individuals. Once validated, real-time feedback could help individuals identify and reproduce conditions associated with greater in-game performance efficiency. Detection of deterioration could instead inform timely rest. Such monitoring could thereby support both optimization of objective task performance and protection from overactivity.

Pupil–heart synchrony could also be used as a readout for testing interventions rather than treated as an established intervention target. Approaches that influence arousal or readiness, including environmental lighting, light exercise, posture, recovery interventions, nutritional strategies, and pre-performance routines, could be tested for their ability to promote constrained physiological reactivity and sustained central–autonomic coordination (49–53). Beyond esports, the same framework could be examined in cognitively demanding digital work and learning in which intense cognitive–sensorimotor demands coexist with limited whole-body movement. Pupil–heart synchrony could form a mechanism-based foundation for adaptive feedback systems that support high-level objective task performance while enabling timely recovery.

## Limitations and future directions

Several limitations point to specific directions for future research. First, the present study identified a session-level association between pupil–heart synchrony and in-game performance efficiency, but it cannot determine whether this coordination plays a causal role or whether the efficient profile represents a reproducible within-person state. Because the physiological clusters were derived from repeated sessions contributed by a modest number of participants, they may also reflect stable differences in arousal regulation, playing strategy, or skill. Repeated within-person studies and interventions that manipulate central arousal or peripheral autonomic activity could establish whether individuals enter and leave this state, whether it can be induced, and whether inducing it improves objective in-game performance while reducing subjective workload.

Second, the selective CRQA pattern provides a testable basis for investigating the mechanisms underlying this state, but pupil diameter and heart rate are indirect physiological indices. The present analyses did not separate tonic from phasic responses or align physiological dynamics to discrete in-game events. Subjective workload was also assessed after active play, so the mediation analysis cannot establish whether reduced workload preceded or resulted from coordinated physiology and superior objective in-game performance. Subjective awareness of efficiency was inferred from subjective flow rather than assessed through direct estimates of performance efficiency or metacognitive accuracy. Event-locked studies combining pupil and cardiac measures with complementary neurophysiological, endocrine, and autonomic indices could determine whether episodes of pupil–heart synchrony coincide with behaviorally relevant events and clarify the temporal relationship among physiological coordination, workload, and objective in-game performance. Direct assessment of *mushin* and *heijoshin* could further test their proposed conceptual correspondence with this physiological organization.

Finally, pupil–heart synchrony was evaluated retrospectively as a biomarker in a single laboratory task and a restricted participant sample rather than prospectively validated. The next critical step is prospective validation across tasks, populations, and naturalistic digital environments. Such studies should establish within-person reliability and classification accuracy and determine whether real-time detection or feedback can improve objective task performance, facilitate reproduction of efficient states, and support timely recovery.

## Conclusion

These findings support the hypothesis that, during game and esports play, central–autonomic coordination reflected in intrapersonal pupil–heart synchrony contributes to an efficient cognitive–sensorimotor performance state that subjective flow fails to capture. Esports competition increased subjective flow without improving objective in-game performance. In contrast, a low-reactivity state with selectively sustained pupil–heart synchrony combined superior objective in-game performance with lower subjective workload, indicating greater in-game performance efficiency. Reduced workload statistically accounted for part of the association between pupil–heart synchrony and objective in-game performance. Subjective– objective miscalibration in modern digital activity extends to efficient cognitive–sensorimotor performance states that individuals may fail to recognize. Pupil–heart synchrony reflects central–autonomic coordination supporting this efficiency. Pupil–heart synchrony may therefore serve as a mechanism-based biomarker for detecting such states beyond subjective awareness, optimizing objective task performance and the timing of recovery in demanding digital environments.

## Materials and Methods

### Participants, ethics approval and consent to participate, and study design

Fourteen healthy young adult men participated in the study. Pupil-diameter data were unavailable for one participant because of a technical recording failure; all analyses reported in this study were therefore restricted to the 13 participants with complete joint pupil and heart-rate data to maintain a common analytical sample across outcomes (mean age, 24.9 ± 1.1 years [mean ± SEM]). The study protocol was approved by the Ethics Review Board of the Institute of Health and Sport Sciences at the University of Tsukuba (approval number: Tai25–90), and all participants provided written informed consent in accordance with the Declaration of Helsinki. Participants completed three 4-min sessions under each of three conditions: passive watching (Watch), playing against a computer-controlled opponent (Game), and competing against another participant (Esports). Watch served as the passive baseline. Further details of participants and experimental procedures are provided in *Appendix*.

### Psychological, in-game performance, and physiological measures

Subjective flow and workload were assessed using the Sports Flow Scale (SFS) and NASA Task Load Index (NASA-TLX), respectively. Objective in-game performance was quantified using knockouts, damage dealt, damage taken, attacking accuracy, and combat efficiency (damage dealt/damage taken). Pupil diameter was recorded at 100 Hz using Tobii Pro Glasses 3 and converted to 1 Hz, whereas heart rate was recorded at 1 Hz using a Polar H10 monitor. For active-play analyses, changes in pupil diameter and heart rate were calculated relative to each participant’s Watch baseline. Detailed measurement and preprocessing procedures are provided in Appendix.

### Clustering of joint pupil–heart reactivity

Unsupervised k-means clustering was applied to 78 active play sessions using z-standardized changes in mean pupil diameter and heart rate relative to the participant-specific Watch baseline. Candidate solutions with k = 2–4 were examined, and k = 3 was retained based on cluster-size balance and physiological interpretability. No psychological, behavioral, or contextual variables were used to define the clusters or select k. Full procedures are provided in Appendix.

### Pupil–heart synchrony analyses

Pupil–heart synchrony was assessed within each participant and active-play session using cross-recurrence quantification analysis (CRQA) of aligned 1-Hz pupil-diameter and heart-rate time series. Recurrence rate was fixed at 5%, with a smaller recurrence radius indicating more tightly constrained joint dynamics. Cross-correlation was used as a linear benchmark. Full procedures are provided in Appendix.

### Statistical analysis

Condition-level comparisons used repeated-measures ANOVA or paired t tests, as appropriate. Session-level analyses used linear mixed-effects models accounting for repeated observations within participants. Mediation analyses examined whether subjective workload statistically accounted for associations between CRQA recurrence radius and objective in-game performance. Full statistical procedures are provided in Appendix and Supplementary Tables.

## Acknowledgement and funding source

This research was supported by Precursory Research for Embryonic Science and Technology (PRESTO) from the Japan Science and Technology Agency (JST) to S.D. (JPMJPR24I8); a Grant-in-Aid for Early-Career Scientists from Japan Society for the Promotion of Science (JSPS) to S.D. (24K18207); a research grant from the Advanced Research Initiative for Human High Performance (ARIHHP), University of Tsukuba to S.D. (2026(I)11); Fusion Oriented Research for disruptive Science and Technology (FOREST) from JST to T.M. (JPMJFR205M); and the Science and Technology Promotion Project in Special Power Source Prefectures (MEXT Subsidy Project) by Ministry of Education, Culture, Sports, Science and Technology (MEXT) to T.M. Funding sources had no involvement in study design; in the collection, analysis and interpretation of data; in the writing of the report; and in the decision to submit the article for publication.

## Data availability

All study data are included in the article and/or Appendix.

## Author contribution

S.D. and T.M. conceived and designed the study. S.D. recruited participants. S.D., T.S., T.Y., Y.T., R.K., and T.M. collected, analyzed, and interpreted the data. S.D. and T.M. drafted the manuscript. T.M. edited and revised the manuscript. All authors approved the manuscript.

## Ethics approval and consent to participate

The study protocol was approved by the Ethics Review Board of the Institute of Health and Sport Sciences at the University of Tsukuba (approval number: Tai25–90), and all participants provided written informed consent in accordance with the Declaration of Helsinki.

## Appendix: Supplementary Methods

### Participant eligibility and sample characterization

Participants were recruited from the local university community and surrounding area. Eligibility criteria were an age of 18–35 years, the capacity to understand the study procedures and provide written informed consent, normal color vision, and availability to attend the laboratory experiment in person. Participant characteristics are summarized in Supplementary Table 1.

All 14 participants completed the experimental protocol. Pupil-diameter data from one participant were unavailable because of a technical recording failure. To maintain a common analytical sample across outcomes, the analyses reported here were restricted to the 13 participants with complete joint pupil and heart-rate data.

### Pre-experiment questionnaire

Following previous studies (1, 2), demographic characteristics, video-game and esports experience, habitual physical activity, and sleep duration were assessed using an online questionnaire administered through Google Forms. Participants received the survey link and study information by email at recruitment and completed the questionnaire by the morning preceding the experiment.

Experience with the game was characterized by self-reported experience with the game title used in the experiment, expressed in months, and average weekday and weekend playing time, expressed in minutes per day. Habitual physical activity was assessed using the validated Japanese version of the International Physical Activity Questionnaire (IPAQ) (3). These questionnaire variables were used to characterize the sample and were not entered as covariates in the primary hypothesis-testing models.

### Experimental setting, game settings, and session order

Experiments were conducted in a climate-controlled laboratory. Ambient temperature was maintained at 24–25 °C, relative humidity at 40–60%, and illuminance at 470–500 lux at eye level. The two participants were seated in the same room at desks arranged in a V-shaped configuration, with each participant viewing an individual display. Participants were instructed to attend to their own display and to refrain from unnecessary verbal communication. Naturally occurring nonverbal behavior, including facial expressions, was not restricted.

Game settings were standardized across sessions. Each participant selected a preferred character before the experiment and used that character throughout the protocol. The stage was fixed to “Battlefield,” all items were disabled, and matches were conducted using a 4-min time-limit rule. The number of knockouts determined the match winner. Each 4-min session was followed by a 3-min questionnaire period.

In the Game condition, computer-opponent difficulty was selected individually during preliminary testing using the game’s nine-level difficulty scale. The selected level was the highest level at which the participant reported being able to play enjoyably while maintaining concentration. The mean selected difficulty was 7.6 ± 0.5. For the Esports condition, participant pairs were formed after preliminary matches confirmed broadly comparable skill levels and the absence of consistently one-sided matches.

Within each pair, participants were assigned to Player A or Player B by the experimenter, and these labels were used to implement the session order. During sessions 1–3, Player A completed three Game sessions while Player B completed three Watch sessions.

During sessions 4–6, the two participants competed against each other in three Esports competition sessions. During sessions 7–9, Player B completed three Game sessions while Player A completed three Watch sessions. This arrangement counterbalanced whether Watch or Game occurred before the Esports condition, although the Esports condition was always performed in the middle block.

### Subjective flow

Subjective flow was assessed using the six-item Sports Flow Scale (SFS), which was developed and validated for the assessment of flow immediately following a sports-related activity (4). The six items assess fusion of action and perception, concentration on the current task, sense of control, loss of self-consciousness, transformation of the sense of time, and enjoyment of the activity as an intrinsically rewarding experience.

Participants rated each item immediately after the preceding 4-min session. Five items were rated from 0 (“not at all”) to 10 (“completely”). For the transformation-of-time item, ratings ranged from 0 (“slower than usual”) through 5 (“unchanged”) to 10 (“faster than usual”). Following the original scoring procedure, this item was transformed as |raw rating − 5| × 2. The SFS total score was calculated by summing the six item scores, producing a possible range of 0– 60, with higher scores indicating stronger subjective flow. The six component scores were also examined separately. The same administration and scoring procedures were used following Watch, Game, and Esports sessions, with instructions referring to the activity experienced during the immediately preceding session.

### Subjective workload

Subjective workload was assessed after each Game and Esports session using the NASA Task Load Index (NASA-TLX) (5). The Japanese-language procedure described by Miyake and

Kumashiro was used (6). The NASA-TLX comprises six dimensions: mental demand, physical demand, temporal demand, own performance, effort, and frustration. Each dimension was rated using a visual analog scale ranging from 0 to 100 (6). Higher values indicated greater demand, effort, or frustration. The own-performance dimension was coded so that higher values represented poorer perceived performance and therefore greater subjective workload.

The primary total workload score used throughout the study was the Adaptive Weighted Workload score (AWWL) (6). For each assessment, the six raw dimension scores were ranked from highest to lowest and assigned weights from 6 to 1, respectively. The AWWL score was calculated as the weighted mean of the six ratings: AWWL = Σ(wᵢ × rᵢ) / Σwᵢ, where rᵢ is the raw rating and wᵢ is its rank-based weight. Higher total scores indicated greater subjective workload. The six dimension scores were additionally analyzed to characterize the source of workload.

### Objective in-game performance data handling

The number of knockouts directly contributed to the match outcome. Damage dealt represented offensive output, damage taken represented defensive cost, and attacking accuracy was treated as the game-reported index of attack success. Combat efficiency integrated offensive output and defensive cost as damage dealt divided by damage taken, with larger values indicating that greater damage was delivered relative to the damage received.

Damage-taken and attacking-accuracy data could not be extracted for one active play session because of a screen-recording failure. Analyses involving these outcomes or combat efficiency were conducted using complete cases. No missing outcome values were imputed.

### Pupil-diameter and heart-rate acquisition, preprocessing, and temporal alignment

Pupil diameter was recorded continuously using an infrared wearable eye tracker sampled at 100 Hz (Tobii Pro Glasses 3; Tobii Technology, Sweden). Calibration was performed before the experimental protocol. Under standardized ambient illumination (470–500 lx), pupil diameter was treated as an index of pupil-linked arousal rather than as a direct measure of activity in a specific brain region.

Pupil-diameter data were processed using the Average eye-selection setting in Tobii Pro Lab, whereby left- and right-eye pupil-diameter values were averaged when both were available and the available eye was used when only one eye was valid. The built-in Noise reduction filter was applied. Samples marked as invalid by the eye-tracking system or affected by blinks were treated as missing, and no interpolation was performed. The remaining valid samples were averaged within non-overlapping 1-s windows to produce a 1-Hz pupil-diameter time series, without applying a predefined minimum valid-sample threshold. Session events were annotated by visual inspection of the recorded video in Tobii Pro Lab, and the corresponding event timestamps were used to identify each 4-min session interval.

Heart rate was recorded using a chest-worn monitor (Polar H10; Polar Electro, Finland), and heart-rate data were obtained from Polar Flow (Polar Electro), the manufacturer-provided software platform, as a 1-Hz time series for joint analysis with the 1-Hz pupil-diameter time series. No additional resampling, interpolation, or artifact correction was applied to the exported heart-rate data.

During the experiment, the experimenter manually recorded the clock time of each session event using a web-based standard time reference. Heart-rate segments were extracted from the Polar Flow recordings based on these recorded clock times and temporally matched to the corresponding session-event timestamps identified in Tobii Pro Lab. The resulting 1-Hz pupil-diameter and heart-rate time series were aligned to the onset of each session, yielding up to 240 paired observations per 4-min session.

### Clustering implementation

Unsupervised k-means clustering was used to identify session-level patterns of joint pupil–heart reactivity (7). The analytical unit was an active play session. The 13 participants with complete joint pupil and heart-rate data each contributed six active play sessions, yielding 78 session-level observations.

The two clustering features were the change in mean pupil diameter and the change in mean heart rate relative to the participant-specific Watch baseline. Both features were standardized across the 78 sessions using z scores before clustering. Clustering was performed using Euclidean distance with randomly initialized cluster centers, a fixed random seed (12345), and a maximum of 100 iterations. Candidate solutions with k = 2, 3, and 4 were examined. A three-cluster solution was retained considering the balance of cluster sizes and the ability to distinguish physiologically interpretable patterns of joint pupil–heart reactivity while retaining sufficient observations within each cluster. Cluster formation and selection of *k* were solely on pupil-diameter and heart-rate reactivity; subjective flow, subjective workload, objective in-game performance, play condition, and win–loss outcome were not used to define the clusters or to select *k*.

Clusters were ordered descriptively according to their centroids and designated as low-, intermediate-, and high-reactivity states. After unsupervised clustering, the clusters were compared with respect to subjective flow, subjective workload, objective in-game performance, cross-correlation measures, and CRQA indices. A cluster was operationally interpreted as an efficient cognitive–sensorimotor performance state only when it combined superior objective in-game performance with lower subjective workload.

### Cross-recurrence quantification analysis

Cross-recurrence quantification analysis (CRQA) was applied to the aligned pupil-diameter and heart-rate time series from each active play session to quantify nonlinear temporal coordination between the two physiological signals (8, 9). Related recurrence-based analysis has been applied to heart-rate synchrony in elite esports teams (10). CRQA was implemented in Python 3.13.9 using PyRQA 8.1.0, NumPy 2.3.5, and SciPy 1.16.3. PyRQA was used for the core cross-recurrence analysis, with additional diagonal- and vertical-line metrics calculated from the recurrence matrices using custom Python routines.

Before CRQA, each 1-Hz pupil and heart-rate time series was linearly detrended and z-standardized within the session. Phase-space reconstruction was performed using time-delay embedding with an embedding dimension of m = 3 and a time delay of τ = 2 samples, corresponding to 2 s at the 1-Hz sampling rate (11, 12). Cross-recurrence matrices were constructed using the Euclidean distance between the reconstructed pupil and heart-rate state vectors. A cross-recurrence point was defined when this distance was less than or equal to the recurrence threshold ε (13).

To enable comparison across sessions, the recurrence threshold was determined separately for each session under a fixed recurrence-rate criterion (14). A bisection algorithm adjusted ε over a prespecified range of 0.02–2.0 for 18 iterations, targeting a cross-recurrence rate of 5% (RR = 0.05). The number of iterations was fixed a priori, and no separate convergence-tolerance criterion was applied. A Theiler window of one sample was applied to exclude recurrences arising from immediate temporal adjacency, and minimum diagonal and vertical line lengths were both set to two samples (lmin = 2 and vmin = 2, respectively) before calculating line-based indices.

The CRQA outcomes were recurrence radius, determinism, average diagonal line length, maximum diagonal line length, diagonal-line entropy, laminarity, average vertical line length, maximum vertical line length, and vertical-line entropy. Because recurrence rate was fixed at 5%, recurrence rate itself was not analyzed as an outcome. The radius required to achieve the fixed recurrence rate was retained as the primary proximity-based index: a smaller radius indicated that the reconstructed pupil and heart trajectories achieved the same recurrence rate within a more tightly constrained joint state space. Maximum diagonal line length indexed the longest episode of sustained similar temporal evolution between the two signals.

In the present study, pupil–heart synchrony refers to the pattern of constrained and sustained pupil–heart coordination jointly characterized by recurrence radius and diagonal-line structure. It does not imply identical signal amplitudes or zero-lag linear correlation.

### Cross-correlation analysis

Cross-correlation analysis was used as a conventional linear benchmark for the CRQA-based assessment of pupil–heart synchrony, following our previous analytical approach (15). Analyses were performed in R version 4.4.0 (R Foundation for Statistical Computing, Vienna, Austria).

The same linearly detrended and within-session z-standardized 1-Hz time series used for CRQA were analyzed. Missing values were interpolated using spline interpolation with the na_interpolation() function in the imputeTS package before analysis. Cross-correlation functions were calculated using the ccf() function in the R stats package across temporal lags from −5 to +5 s. For each session, the largest positive cross-correlation coefficient within this interval was retained as the maximum cross-correlation. The lag at which this maximum occurred was also identified, and its absolute value was retained as an index of temporal alignment irrespective of direction; smaller absolute lag values indicated closer linear temporal correspondence.

### Sample-size calculation

An a priori power analysis was conducted in G*Power 3.1.9.7 for a two-tailed paired comparison, assuming Cohen’s dz = 0.80, α = 0.05, and 80% power. This analysis indicated that at least 12 participants were required. Fourteen participants were recruited to allow for potential data loss.

All 14 participants completed the experimental protocol. Because pupil-diameter data from one participant were unavailable owing to a technical recording failure, all analyses reported in this study were restricted to the 13 participants with complete joint pupil and heart-rate data to maintain a common analytical sample across outcomes. These 13 participants contributed 78 active play sessions to the session-level analyses. The clustering and mediation analyses were exploratory session-level analyses with repeated observations nested within participants; the original participant-level power calculation was not intended as a confirmatory power analysis for these models.

### Detailed statistical analysis

Statistical analyses were performed using GraphPad Prism 10.6.0 (GraphPad Software, San Diego, CA, USA), StataNow 19 SE (StataCorp LLC, College Station, TX, USA), R 4.4.0, and Python 3.13.9. All tests were two-sided, and statistical significance was defined as p < 0.05.

Descriptive data are presented as mean ± SEM unless otherwise specified. No missing outcome values were imputed for inferential statistical analyses.

For condition-level comparisons, measurements from the three sessions within each condition were averaged for each participant. SFS total and component scores and absolute pupil-diameter and heart-rate values were compared across Watch, Game, and Esports using one-way repeated-measures ANOVA. The Geisser–Greenhouse correction was applied when sphericity was not assumed. Significant omnibus effects were followed by Bonferroni-corrected paired comparisons. Partial eta squared (η²p) was reported for omnibus effects.

NASA-TLX scores, objective in-game performance outcomes, and pupil- and heart-rate changes from the Watch baseline were compared between Game and Esports using paired t tests based on participant-level condition means. Cohen’s dz was calculated using the mean within-participant difference divided by the standard deviation of the within-participant differences.

The validity of the objective in-game performance outcomes was examined by comparing winning and losing observations using linear mixed-effects models. Win–loss status was entered as a fixed effect, and participant identity and match identity were included as crossed random intercepts. Model-derived estimated marginal means and unstandardized coefficients are reported.

Associations between subjective flow and objective in-game performance were examined at the session level using linear mixed-effects models. SFS total score was entered as a fixed effect, and participant identity was included as a random intercept. The prespecified outcomes were the number of knockouts and combat efficiency. P values were adjusted across these two outcomes using the Benjamini–Hochberg false-discovery-rate procedure.

For comparisons across the physiological clusters, each active play session was treated as an analytical unit. Linear mixed-effects models included cluster as a fixed effect and participant identity as a random intercept. Outcomes were subjective flow, subjective workload, objective in-game performance, cross-correlation indices, and CRQA indices. Significant cluster effects were followed by Bonferroni-corrected contrasts of model-derived estimated marginal means. Unless otherwise specified, cluster figures present estimated marginal means ± SEM.

The association between gameplay context (Game vs. Esports) and physiological cluster membership was examined using multinomial logistic regression, with physiological cluster specified as the outcome, gameplay context as the predictor, and standard errors clustered by participant. Win–loss outcomes across physiological clusters were examined using mixed-effects logistic regression, with cluster specified as a fixed effect and participant identity specified as a random intercept. Overall associations were evaluated using Wald chi-square tests. Mediation analyses examined whether subjective workload statistically accounted for part of the association between pupil–heart synchrony and objective in-game performance.

CRQA recurrence radius was entered as the predictor, NASA-TLX total score as the mediator, and the number of knockouts, combat efficiency, damage dealt, or damage taken as the outcome. A smaller radius represented more tightly constrained pupil–heart dynamics. The indirect effect was calculated as the product of the predictor-to-mediator and mediator-to-outcome coefficients.

Unstandardized coefficients (b) and 95% confidence intervals are reported, and an indirect effect was considered statistically significant when its confidence interval did not include zero. Because these models were based on observational associations within the experimental sessions, mediation was interpreted statistically rather than causally.

Model assumptions were evaluated using residual and quantile–quantile plots. Exact test statistics, effect sizes, confidence intervals, and adjusted post hoc comparisons are provided in the Supplementary Tables.

## Appendix: Figures

**Supplementary Figure 1.**
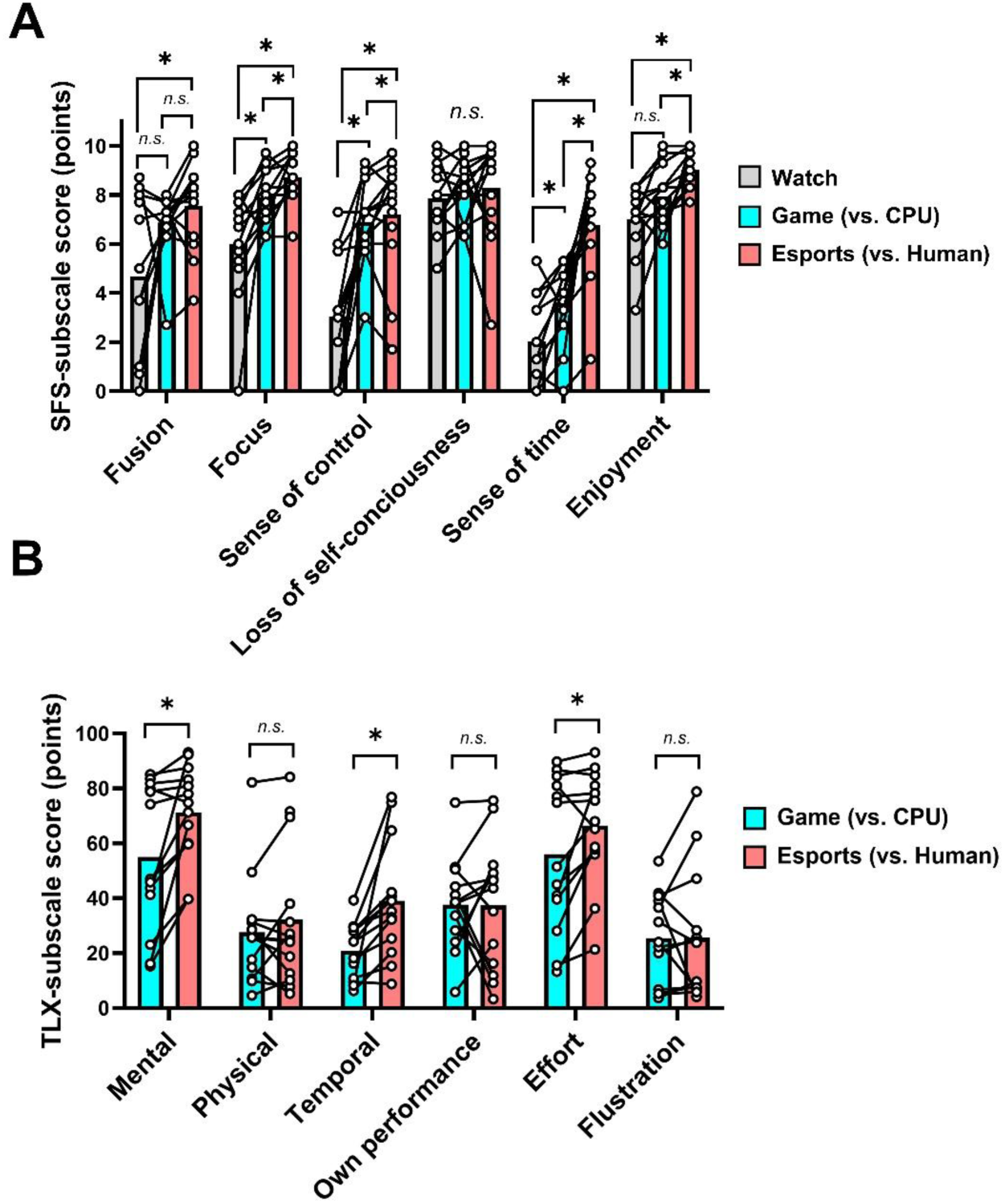
Sports Flow Scale and NASA-TLX subscale scores under the Watch, Game, and Esports conditions. **A.** SFS-subscale scores (“fusion of action and perception”, “focus on current task”, “sense of control”, “loss of self-consciousness”, “transformation of the sense of time”, and “level of enjoyment in the activity”). **B.** NASA Task Load Index (NASA-TLX) subscale scores (“mental demand”, “physical demand”, “temporal demand”, “own performance”, “sense of effort”, and “frustration”). Bars represent group means, and plots represent individual participants. Differences among conditions were assessed using one-way repeated-measures ANOVA or paired t tests. Post hoc comparisons following one-way repeated-measures ANOVA were performed using Bonferroni’s multiple-comparison test. *p < 0.05. n.s., not significant. Details of the statistical analysis are provided in Supplementary Tables 2 and 3.

**Supplementary Figure 2.**
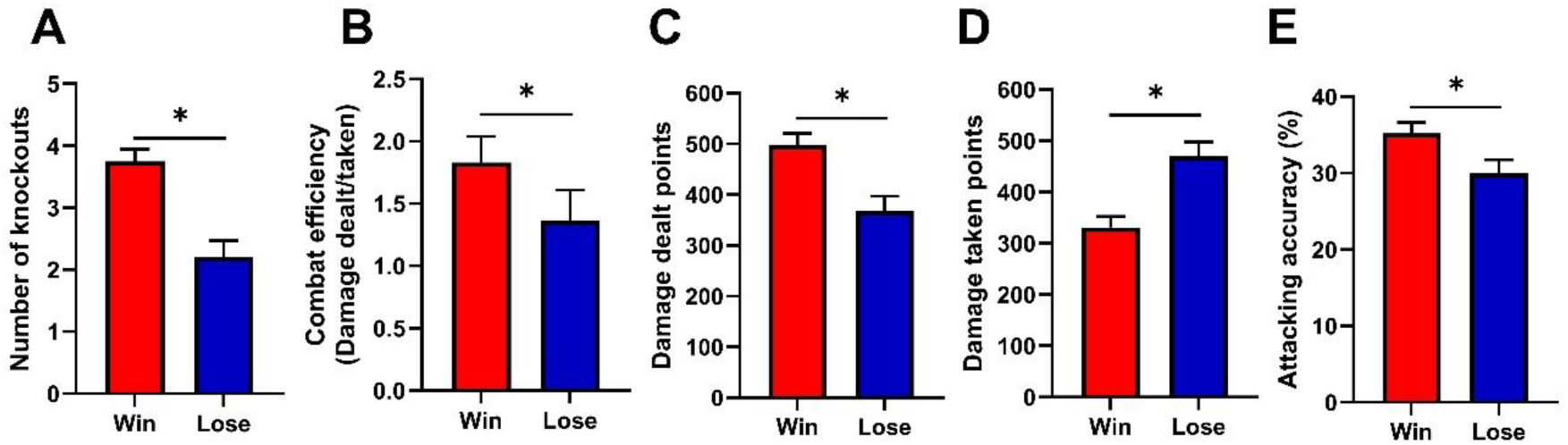
Validation of objective in-game performance metrics by win–loss outcomes. **A.** Number of knockouts. **B.** Combat efficiency, calculated as damage dealt divided by damage taken. **C.** Damage dealt points. **D.** Damage taken points. **E.** Attacking accuracy. Estimated marginal means ± SEM were derived from linear mixed-effects models including win–loss status as a fixed effect and participant identity and match identity as crossed random intercepts. *p < 0.05.

**Supplementary Figure 3.**
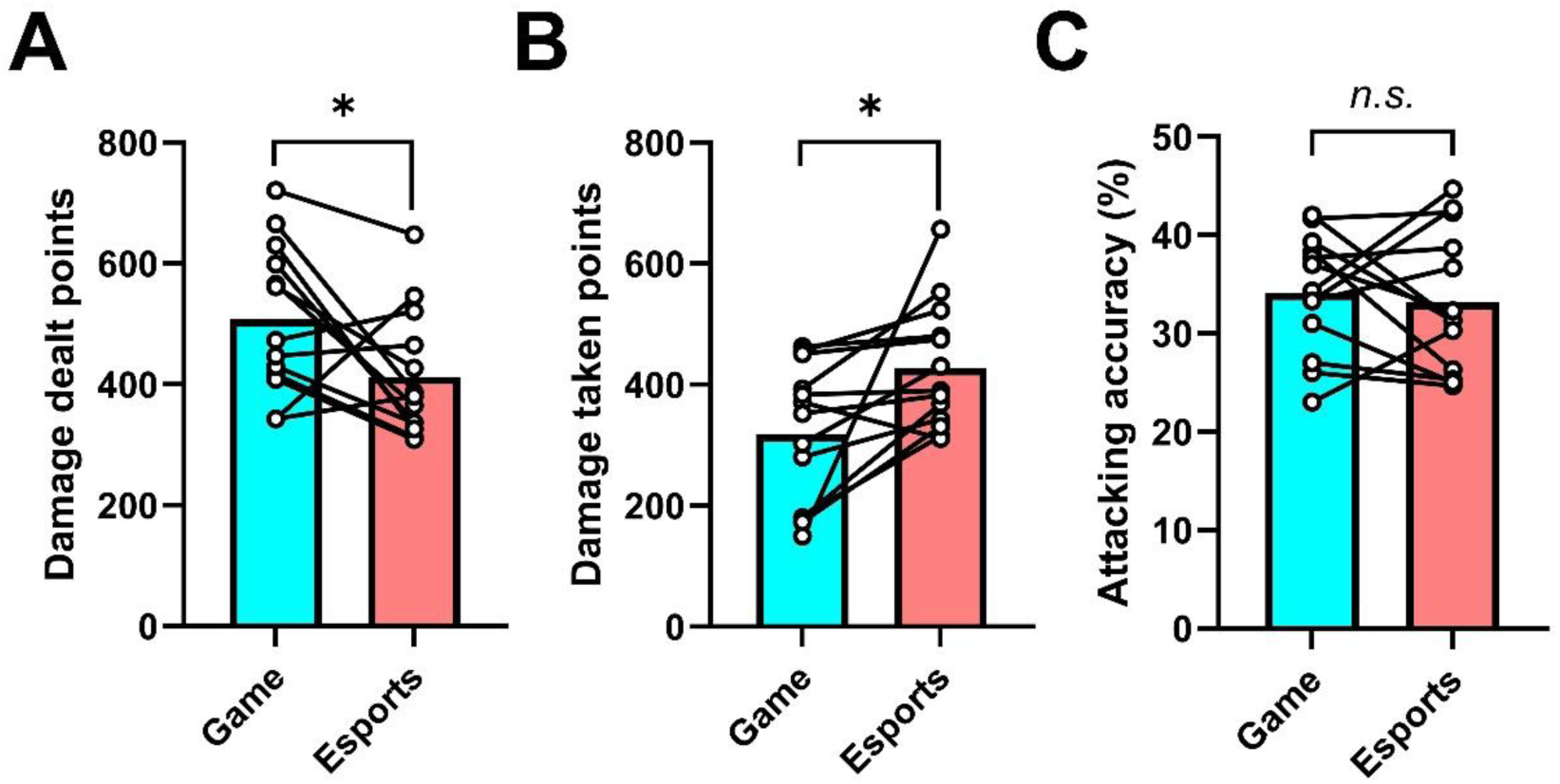
Objective in-game performance under the Game and Esports conditions not shown in Figure 1. **A.** Damage dealt; **B.** Damage taken; **C.** Attacking accuracy. Bars represent group means, and plots represent individual participants. Statistical comparisons were performed using paired t tests. *p < 0.05. n.s., not significant. Details of the statistical analysis are provided in Supplementary Table 2.

**Supplementary Figure 4.**
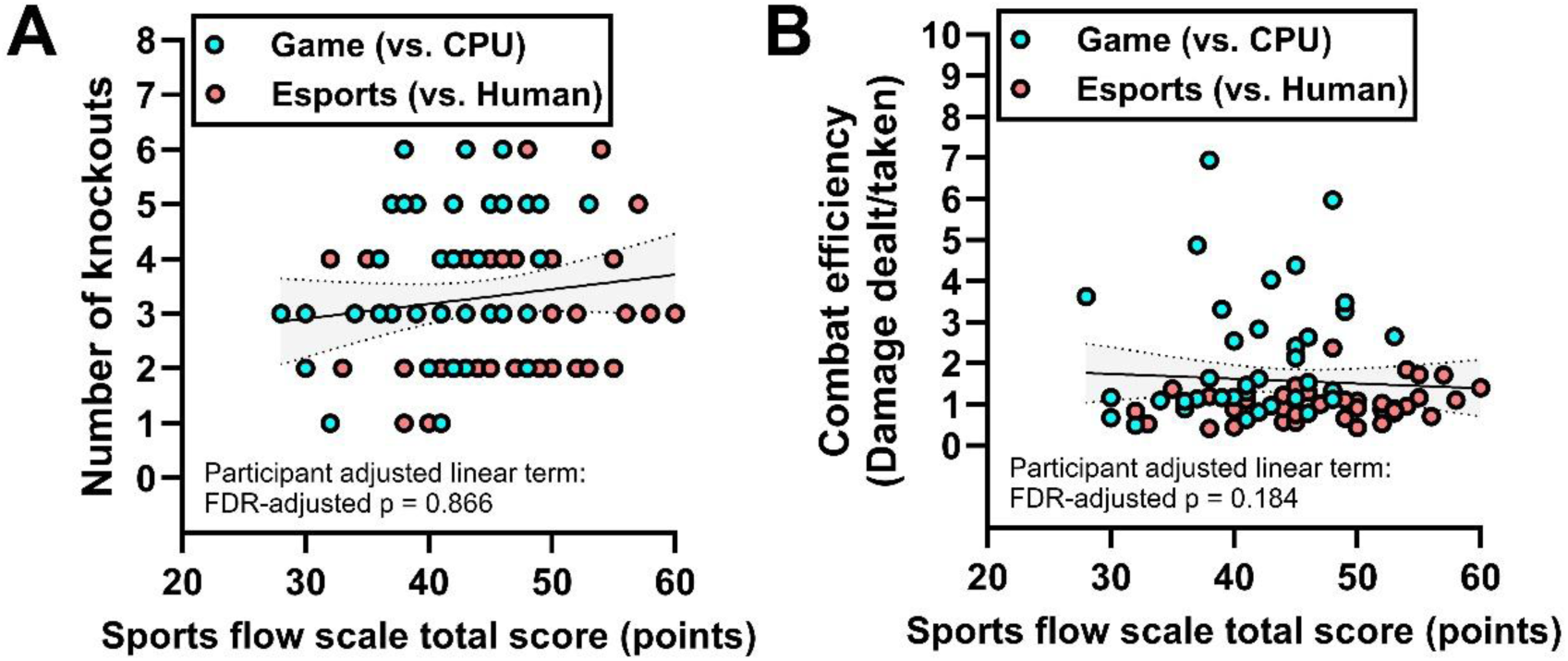
Subjective flow is not associated with objective in-game performance. **A–B**. Relationships between Sports Flow Scale total score and objective in-game performance outcomes, including number of knockouts (**A**) and combat efficiency (**B**). Data points represent active play sessions and are colored by play condition. Lines represent linear fits for visualization, with dotted lines indicating 95% confidence intervals. Statistical significance of linear terms was tested using linear mixed-effects models with participant ID as a random effect. P-values were adjusted using the false discovery rate (FDR) method across the two objective in-game performance outcomes. No significant associations were observed, indicating that subjective flow did not reliably explain high in-game performance efficiency.

**Supplementary Figure 5.**
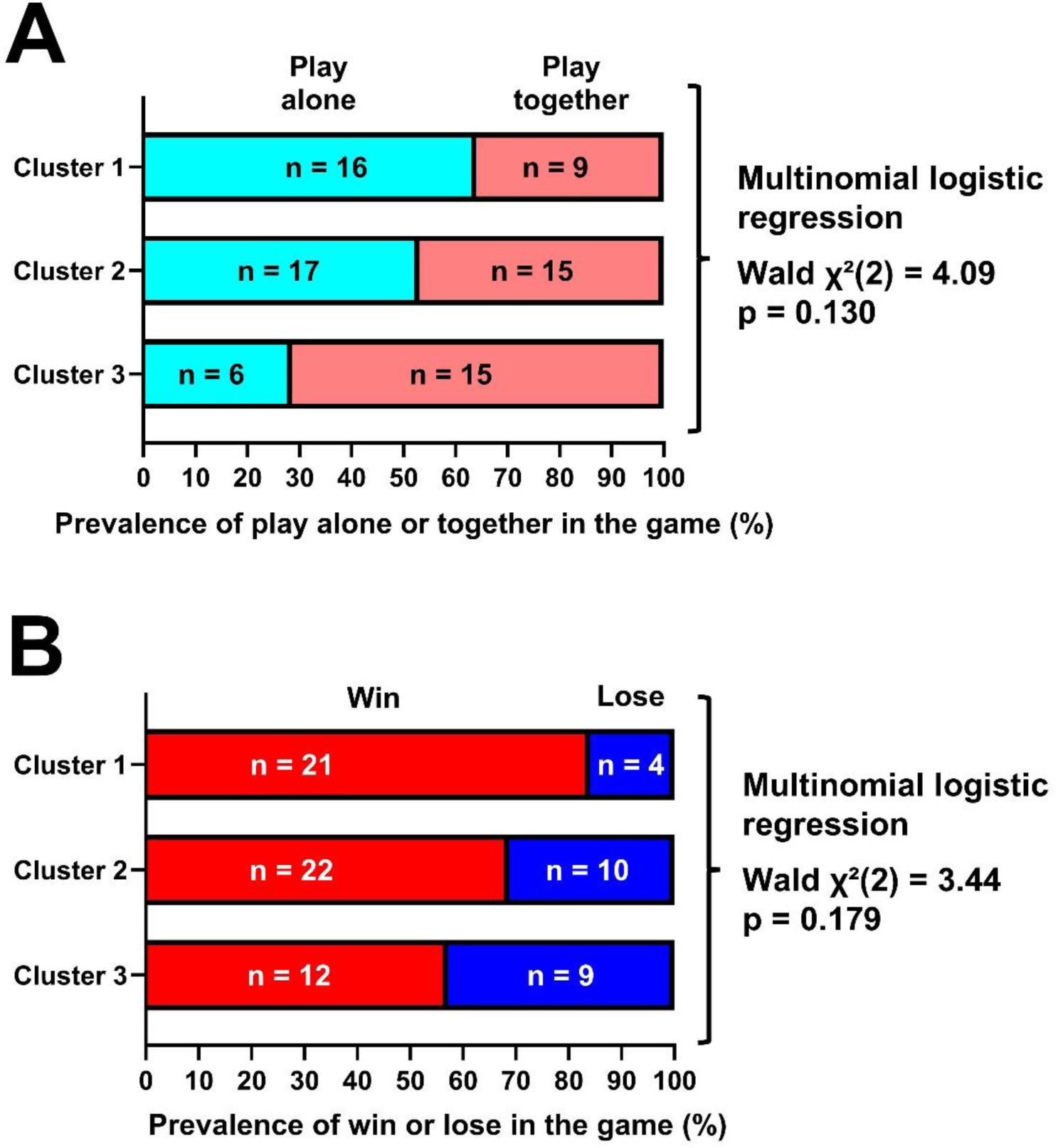
Distributions of play condition and win–loss outcomes did not differ significantly across physiological reactivity clusters. **A.** Distribution of Game and Esports sessions within each physiological cluster identified by k-means clustering of pupil-diameter and heart-rate reactivity. The association between gameplay context and physiological cluster membership was examined using multinomial logistic regression with standard errors clustered by participant. **B.** Distribution of win and loss outcomes within each physiological cluster. Win–loss outcomes were examined using mixed-effects logistic regression, with physiological cluster specified as a fixed effect and participant identity specified as a random intercept. Bars show the observed proportion of gameplay sessions in each category, with the corresponding number of sessions indicated within each bar. Overall associations were evaluated using Wald chi-square tests, with p values shown in each panel.

**Supplementary Figure 6.**
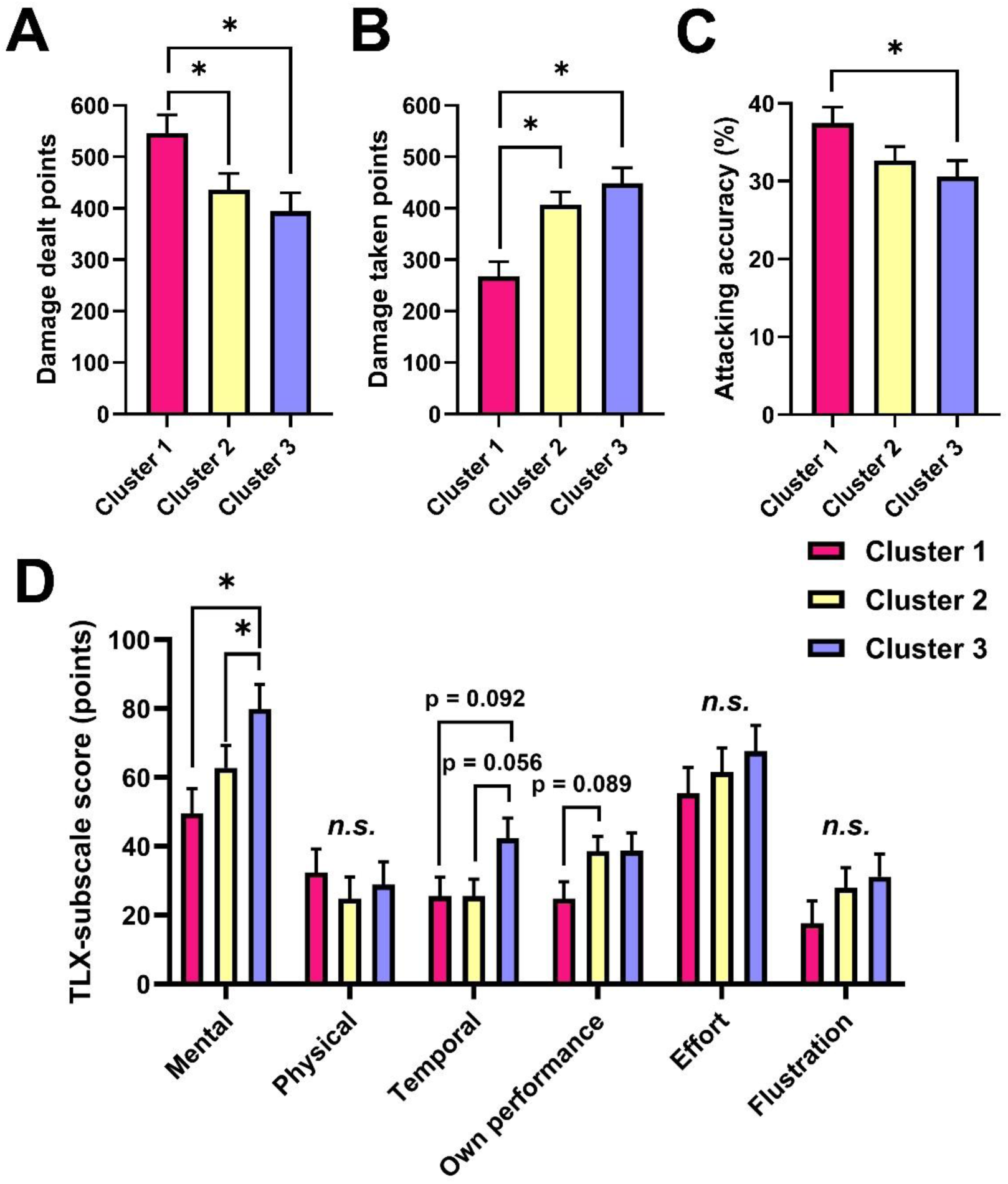
Additional objective in-game performance and subjective workload profiles across physiological clusters. **A.** Damage dealt, reflecting offensive output; **B.** damage taken, reflecting defensive cost; **C.** attacking accuracy, reflecting the precision of offensive actions; and **D.** NASA-TLX subscale scores across the three physiological clusters. Bars show estimated marginal means ± SEM from linear mixed-effects models (LMMs) with participant ID as a random intercept. Post hoc comparisons were conducted using Bonferroni-corrected contrasts of estimated marginal means. *p < 0.05, n.s., not significant. Details of the statistical analysis are provided in Supplementary Tables 4 and 6.

**Supplementary Figure 7.**
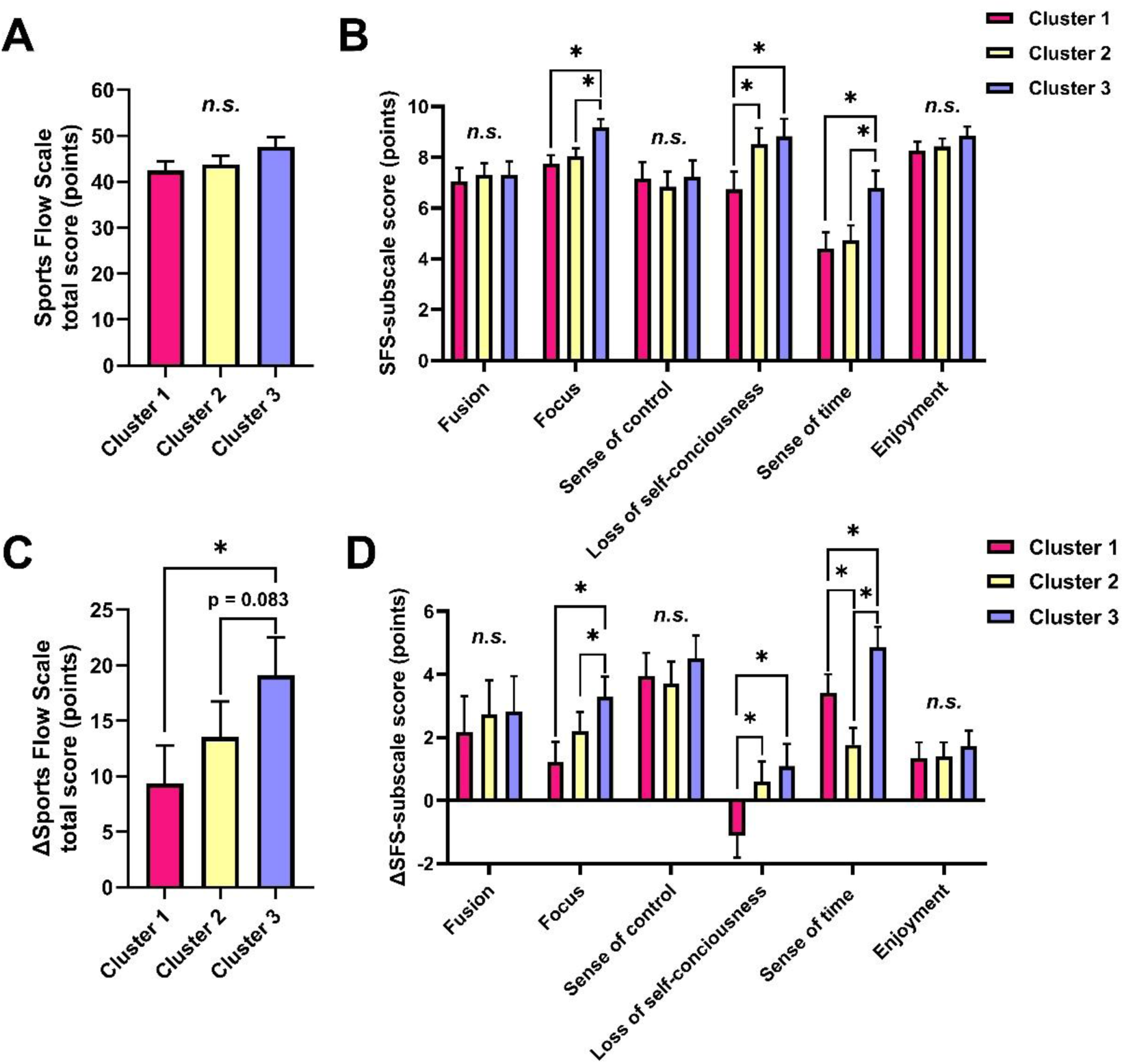
Absolute and change scores of subjective flow across physiological clusters. Absolute Sports Flow Scale (SFS) total and subscale scores (“fusion of action and perception”, “focus on current task”, “sense of control”, “loss of self-consciousness”, “transformation of the sense of time”, and “level of enjoyment in the activity”), together with changes in SFS scores from the watching baseline, are shown across the three physiological clusters. The upper panels (**A–B**) show absolute SFS scores, whereas the lower panels (**C–D**) show ΔSFS scores relative to the watching condition. The ΔSFS total score is reproduced from Figure 2E to facilitate comparison with the corresponding subscale changes. Bars indicate mean values, and error bars indicate SEM. *p < 0.05, n.s., not significant. Details of the statistical analysis are provided in Supplementary Tables 7 and 8.

**Supplementary Figure 8.**
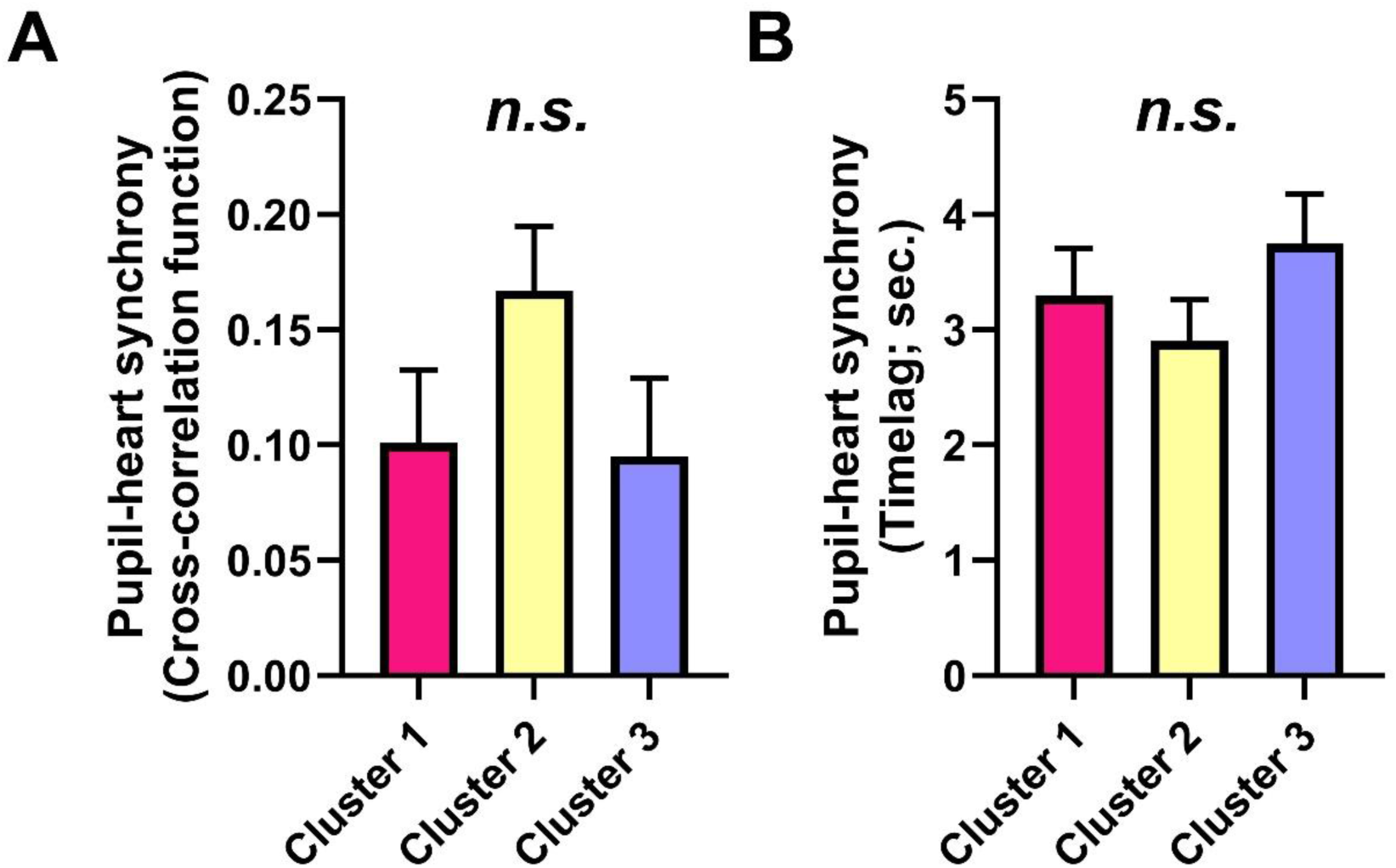
Cross-correlation metrics of pupil–heart dynamics across clusters. **A.** Similarity of pupil–heart dynamics (cross-correlation function). **B**. Time lag of pupil–heart dynamics. Bars show estimated marginal means ± SEM from linear mixed-effects models (LMMs) with participant ID as a random intercept. Post hoc comparisons were conducted using Bonferroni-corrected contrasts of estimated marginal means. n.s., not significant. Details of the statistical analysis are provided in Supplementary Table 9.

**Supplementary Figure 9.**
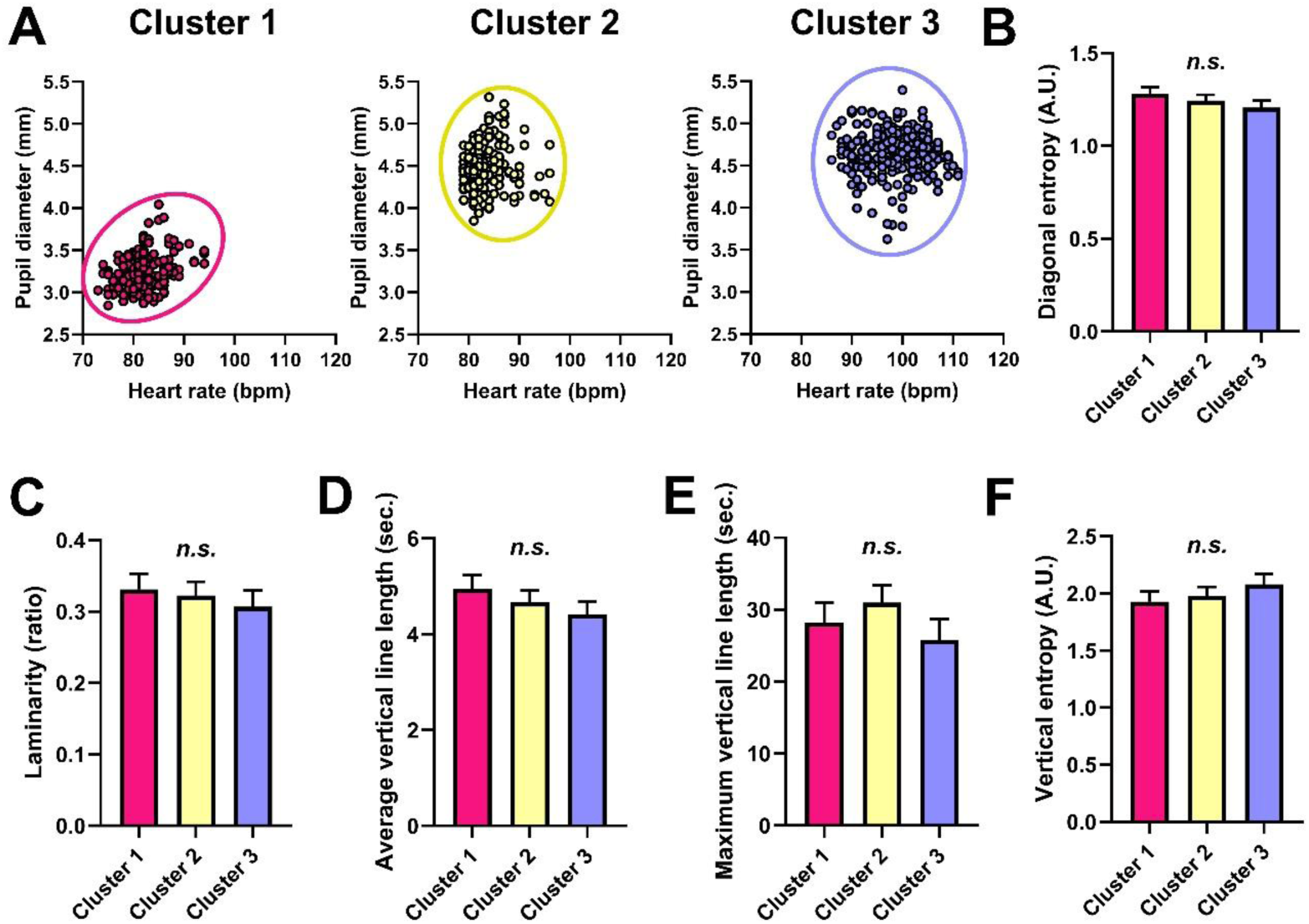
State-space distribution of pupil–heart dynamics and additional cross-recurrence quantification analysis (CRQA) indices across clusters. A. Scatter plots of pupil diameter and heart rate for each cluster. Each point represents a time point in the pupil–heart state space, and ellipses indicate the distribution of data points within each cluster. Clusters were defined based on two-dimensional k-means clustering (k = 3) of changes in mean pupil diameter and heart rate relative to the watching condition (Figure 2A). B–F. Additional CRQA indices across clusters. The following indices are derived from recurrence plot structures, as illustrated in Figure 3A. B. Diagonal entropy, reflecting the complexity of diagonal line length distributions. C. Laminarity, representing the proportion of recurrent points forming vertical line structures. D. Average vertical line length, indicating the mean duration of laminar (stable) states. E. Maximum vertical line length (MVL), representing the longest duration of laminar states. F. Vertical entropy, reflecting the complexity of vertical line length distributions. Bars show estimated marginal means ± SEM from linear mixed-effects models (LMMs) with participant ID as a random intercept. Post hoc comparisons were conducted using Bonferroni-corrected contrasts of estimated marginal means. n.s., not significant. Details of the statistical analysis are provided in Supplementary Table 10.

**Supplementary Figure 10.**
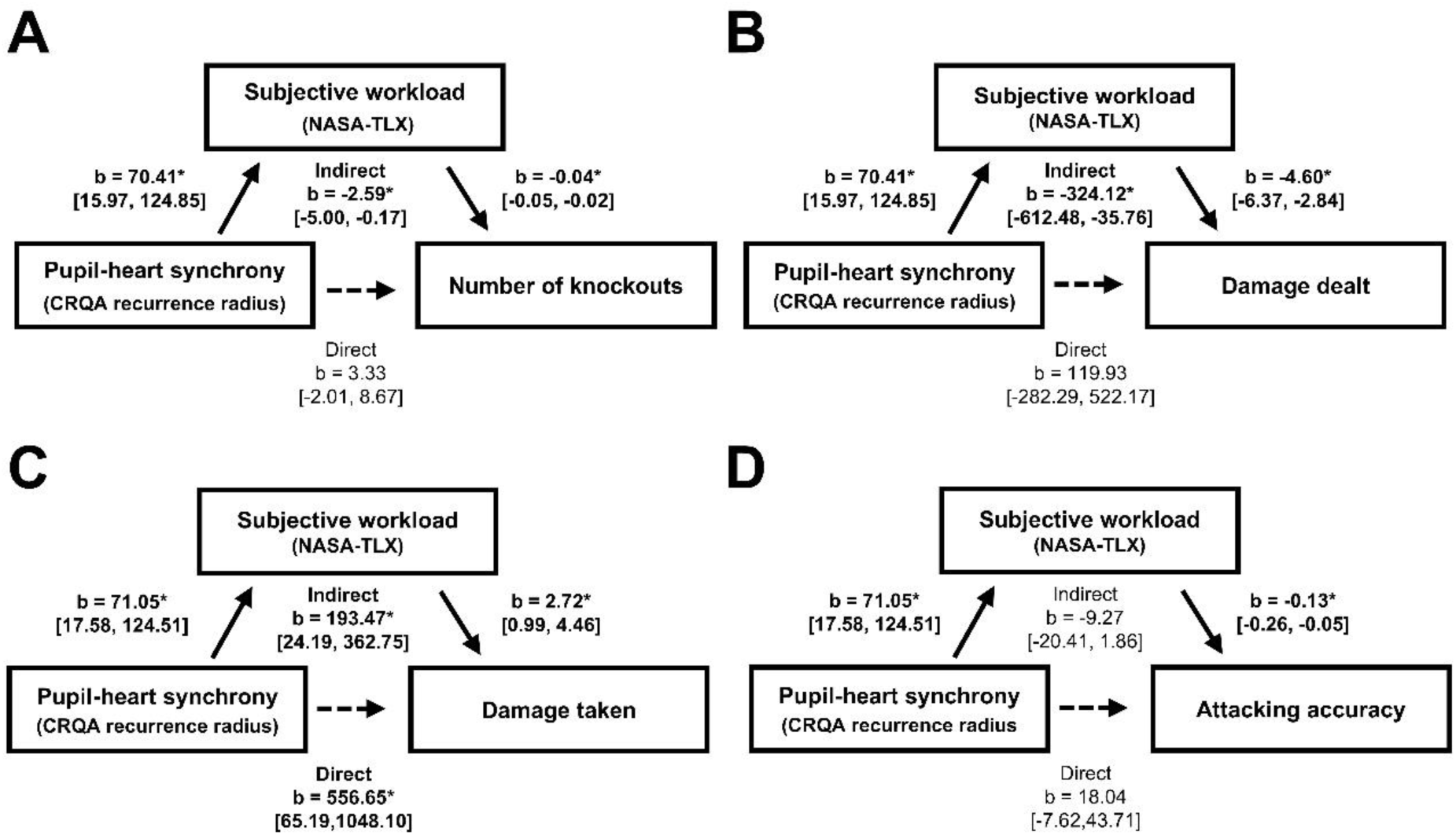
Mediation analysis of pupil–heart synchrony, subjective workload, and objective in-game performance. CRQA recurrence radius was entered as the predictor, NASA Task Load Index (NASA-TLX) total score as the mediator, and number of knockouts (A), damage dealt (B), damage taken (C), or attacking accuracy (D) as the outcome. A smaller recurrence radius indicates more tightly constrained joint pupil–heart dynamics under the fixed recurrence-rate criterion. Indirect effects were calculated as the product of the predictor-to-mediator and mediator-to-outcome coefficients. Values are presented as unstandardized coefficients (b) with 95% confidence intervals. p < 0.05. Detailed statistical results are provided in Supplementary Table 11.

## Appendix: Tables

**Supplementary Table 1.**
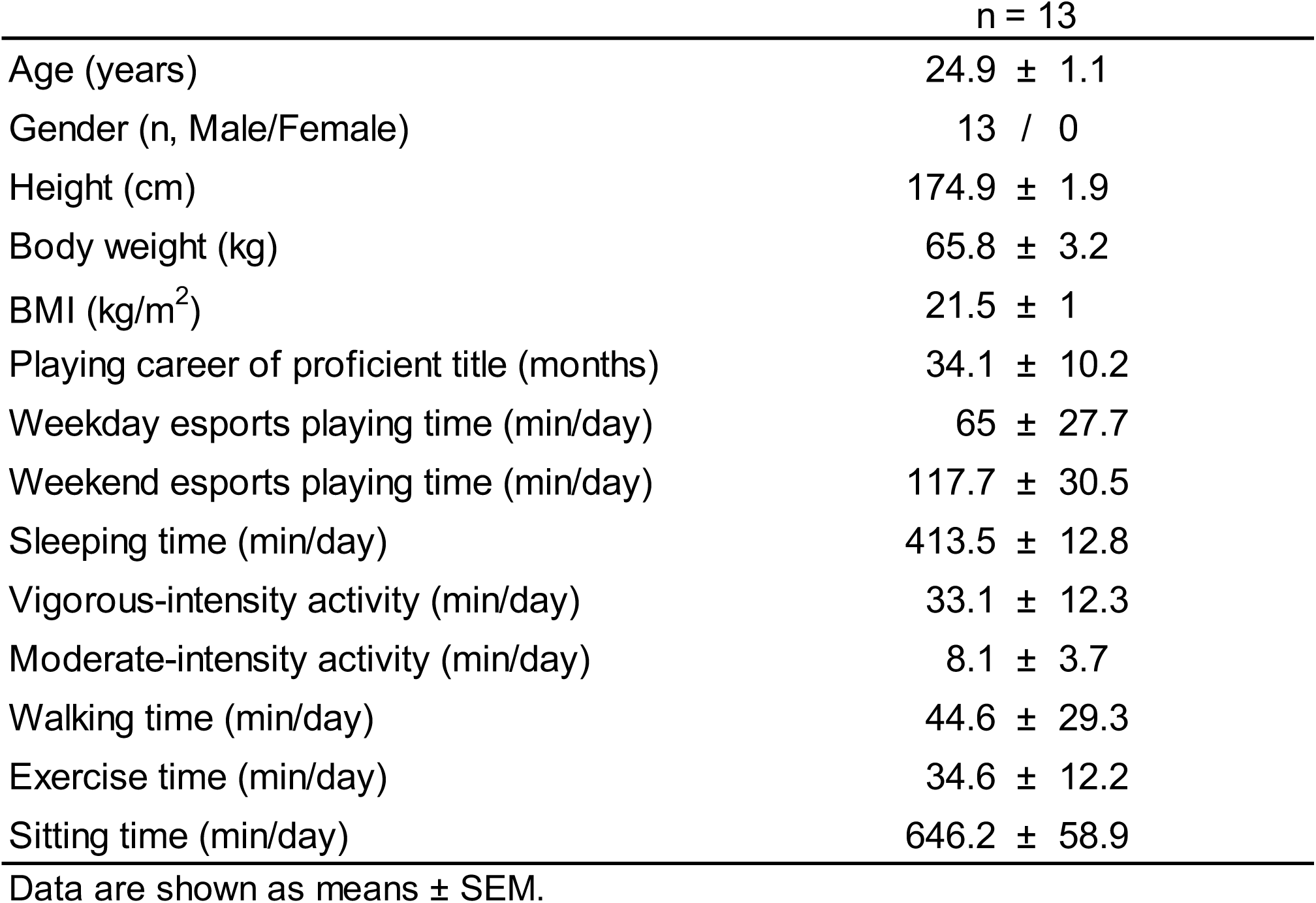
Participant characteristics.

**Supplementary Table 2.** Summary of statistical comparisons among Watch, Game, and Esports conditions for all variables.

| Outcome | F-value or t-value | P-value | Effect size |
| --- | --- | --- | --- |
| Sports Flow Scale scores across Watch, Game, and Esports (Figure 1C, Supplementary Figure 1A) |  |  |  |
| Sports Flow Scale total scores | $F(2,24) = 21.12$ | $< 0.001$ | partial $\eta^2 = 0.637$ |
| Fusion of action and perception | $F(2,24) = 5.35$ | 0.012 | partial $\eta^2 = 0.308$ |
| Focus on current task | $F(2,24) = 13.36$ | $< 0.001$ | partial $\eta^2 = 0.527$ |
| Sense of control | $F(2,24) = 28.47$ | $< 0.001$ | partial $\eta^2 = 0.704$ |
| Loss of self-consciousness | $F(2,24) = 0.29$ | 0.749 | partial $\eta^2 = 0.023$ |
| Transformation of sense of time | $F(2,24) = 33.09$ | $< 0.001$ | partial $\eta^2 = 0.734$ |
| Level of enjoyment in the activity | $F(2,24) = 13.86$ | $< 0.001$ | partial $\eta^2 = 0.536$ |
| NASA-TLX scores across Watch, Game, and Esports (Supplementary Figure 1B) |  |  |  |
| NASA-TLX total scores | $t(12) = 2.87$ | 0.014 | Cohen's $d_z = 0.796$ |
| Mental demand | $t(12) = 3.28$ | 0.006 | Cohen's $d_z = 0.910$ |
| Physical demand | $t(12) = 0.52$ | 0.615 | Cohen's $d_z = 0.143$ |
| Temporal demand | $t(12) = 3.57$ | 0.004 | Cohen's $d_z = 0.993$ |
| Own performance | $t(12) = 0.04$ | 0.969 | Cohen's $d_z = 0.011$ |
| Effort | $t(12) = 2.52$ | 0.027 | Cohen's $d_z = 0.699$ |
| Frustration | $t(12) = 0.04$ | 0.966 | Cohen's $d_z = 0.012$ |
| Game performances across Game and Esports (Figure 1E,F and Supplementary Figure 3A-C) |  |  |  |
| Number of knockouts | $t(12) = 1.52$ | 0.155 | Cohen's $d_z = 0.421$ |
| Combat efficiency | $t(12) = 2.77$ | 0.017 | Cohen's $d_z = 0.768$ |
| Damage dealt points | $t(12) = 2.35$ | 0.037 | Cohen's $d_z = 0.652$ |
| Damage taken points | $t(12) = 2.82$ | 0.016 | Cohen's $d_z = 0.780$ |
| Attacking accuracy | $t(12) = 0.47$ | 0.648 | Cohen's $d_z = 0.133$ |
| Physiological arousal indices across Watch, Game, and Esports (Figure 1G-J) |  |  |  |
| Mean pupil diameter | $F(2,24) = 32.35$ | $< 0.001$ | partial $\eta^2 = 0.729$ |
| Mean heart rate | $F(2,24) = 10.12$ | $< 0.001$ | partial $\eta^2 = 0.457$ |
| ΔMean pupil diameter from Watch condition | $t(12) = 2.35$ | 0.037 | Cohen's $d_z = 0.651$ |
| ΔMean heart rate from Watch condition | $t(12) = 3.14$ | 0.009 | Cohen's $d_z = 0.871$ |

Watch, passive viewing of the game; Game, play against a computer-controlled opponent; Esports, competition against a human opponent. Data are from 13 participants. Outcomes assessed across all three conditions were analyzed using repeated-measures ANOVA, whereas outcomes assessed during active play were compared between Game and Esports using two-sided paired $t$ tests.

**Supplementary Table 3.** Results of Bonferroni post-hoc tests among Watch, Game, and Esports conditions for Sports Flow Scale and physiological arousal variables.

| Outcome | Comparison | Mean diff<br>(absolute values) | Adjusted p-value | Effect size<br>(Cohen's dz) |
| --- | --- | --- | --- | --- |
| Sports Flow Scale (SFS) |  |  |  |  |
| SFS total scores (Figure 1C) | Watch vs Game | 10.92 | 0.003 | 1.211 |
|  | Watch vs Esports | 17.08 | < 0.001 | 1.395 |
|  | Game vs Esports | 6.16 | 0.019 | 0.918 |
| SFS subscales (Supplementary Figure 1A) |  |  |  |  |
| Fusion of action and perception | Watch vs Game | 2.19 | 0.129 | 0.545 |
|  | Watch vs Esports | 2.87 | 0.030 | 0.742 |
|  | Game vs Esports | 0.68 | 0.174 | 0.533 |
| Focus on current task | Watch vs Game | 1.92 | 0.014 | 0.869 |
|  | Watch vs Esports | 2.72 | 0.002 | 1.161 |
|  | Game vs Esports | 0.81 | 0.048 | 0.785 |
| Sense of control | Watch vs Game | 3.84 | < 0.001 | 1.471 |
|  | Watch vs Esports | 4.16 | < 0.001 | 1.747 |
|  | Game vs Esports | 0.32 | > 0.999 | 0.218 |
| Loss of self-consciousness | Watch vs Game | 0.29 | > 0.999 | 0.207 |
|  | Watch vs Esports | 0.42 | > 0.999 | 0.170 |
|  | Game vs Esports | 0.12 | > 0.999 | 0.060 |
| Transformation of sense of time | Watch vs Game | 1.55 | 0.008 | 0.938 |
|  | Watch vs Esports | 4.73 | < 0.001 | 1.856 |
|  | Game vs Esports | 3.18 | 0.002 | 1.501 |
| Level of enjoyment in the activity | Watch vs Game | 0.98 | 0.053 | 0.669 |
|  | Watch vs Esports | 2.03 | < 0.001 | 1.270 |
|  | Game vs Esports | 1.06 | 0.009 | 0.962 |
| Physiological arousal indices (Figure 1G, I) |  |  |  |  |
| Mean pupil diameter | Watch vs Game | 0.54 | < 0.001 | 1.515 |
|  | Watch vs Esports | 0.66 | < 0.001 | 1.847 |
|  | Game vs Esports | 0.12 | 0.139 | 0.573 |
| Mean heart rate | Watch vs Game | 4.21 | 0.439 | 0.537 |
|  | Watch vs Esports | 12.28 | 0.083 | 0.992 |
|  | Game vs Esports | 8.08 | 0.264 | 0.871 |

Watch, passive viewing of the game; Game, play against a computer-controlled opponent; Esports, competition against a human opponent. Data are from 13 participants. Outcomes assessed across all three conditions were analyzed using repeated-measures ANOVA. Multiplicity-adjusted post hoc comparisons were performed where applicable.

**Supplementary Table 4.** Regression coefficients, 95% confidence intervals, and P-values from linear mixed-effects models comparing clusters on the in-game performance (Figure 2B-C; Supplementary Figure 6A-C).

| Outcome | Comparison | Adjusted p-value | b (95 % CI) |
| --- | --- | --- | --- |
| Number of knockouts | Cluster 1 vs Cluster 2 | 0.009 | -1.25 (-2.26, -0.24) |
|  | Cluster 1 vs Cluster 3 | 0.044 | -1.11 (-2.20, -0.02) |
|  | Cluster 2 vs Cluster 3 | > 0.999 | 0.14 (-0.84, 1.12) |
| Combat efficiency | Cluster 1 vs Cluster 2 | 0.001 | -1.17 (-1.96, -0.39) |
|  | Cluster 1 vs Cluster 3 | < 0.001 | -1.46 (-2.33, -0.60) |
|  | Cluster 2 vs Cluster 3 | > 0.999 | -0.29 (-1.09, 0.52) |
| Damage dealt points | Cluster 1 vs Cluster 2 | 0.029 | -109.97 (-211.68, -8.25) |
|  | Cluster 1 vs Cluster 3 | 0.003 | -152.02 (-261.66, -42.37) |
|  | Cluster 2 vs Cluster 3 | 0.907 | -42.05 (-139.66, 55.56) |
| Damage taken points | Cluster 1 vs Cluster 2 | < 0.001 | 139.41 (52.45, 226.37) |
|  | Cluster 1 vs Cluster 3 | < 0.001 | 181.28 (86.11, 276.44) |
|  | Cluster 2 vs Cluster 3 | 0.748 | 41.86 (-45.12, 128.8) |
| Attacking accuracy | Cluster 1 vs Cluster 2 | 0.128 | -4.83 (-10.54, 0.87) |
|  | Cluster 1 vs Cluster 3 | 0.021 | -6.92 (-13.06, -0.78) |
|  | Cluster 2 vs Cluster 3 | > 0.999 | -2.08 (-7.52, 3.34) |

Analyses included 78 active play sessions from 13 participants. b represents the estimated difference in outcome (second cluster minus first cluster) derived from linear mixed-effects models adjusted for repeated measures within participants. CI: confidence interval.

**Supplementary Table 5.** Sensitivity analyses of cluster differences in in-game performance after adjustment for play condition.

| Outcome | Comparison | Adjusted p-value | b (95 % CI) |
| --- | --- | --- | --- |
| Number of knockouts | Cluster 1 vs Cluster 2 | 0.022 | -1.12 (-2.12, -0.12) |
|  | Cluster 1 vs Cluster 3 | 0.359 | -0.75 (-1.90, 0.40) |
|  | Cluster 2 vs Cluster 3 | > 0.999 | 0.37 (-0.63, 1.37) |
| Combat efficiency | Cluster 1 vs Cluster 2 | 0.004 | -1.05 (-1.83, -0.28) |
|  | Cluster 1 vs Cluster 3 | 0.036 | -0.95 (-1.86, -0.05) |
|  | Cluster 2 vs Cluster 3 | > 0.999 | 0.10 (-0.70, 0.91) |
| Damage dealt points | Cluster 1 vs Cluster 2 | 0.097 | -86.19 (-182.63, 10.25) |
|  | Cluster 1 vs Cluster 3 | 0.158 | -90.17 (-201.65, 21.30) |
|  | Cluster 2 vs Cluster 3 | > 0.999 | -3.98 (-99.18, 91.21) |
| Damage taken points | Cluster 1 vs Cluster 2 | 0.001 | 126.46 (40.30, 212.62) |
|  | Cluster 1 vs Cluster 3 | 0.013 | 119.37 (19.11, 219.64) |
|  | Cluster 2 vs Cluster 3 | > 0.999 | -7.09 (-93.49, 79.31) |
| Attacking accuracy | Cluster 1 vs Cluster 2 | 0.114 | -5.03 (-10.85, 0.78) |
|  | Cluster 1 vs Cluster 3 | 0.026 | -7.42 (-14.18, -0.65) |
|  | Cluster 2 vs Cluster 3 | 0.95 | -2.38 (-8.08, 3.31) |

Analyses included 78 active play sessions from 13 participants, except for damage taken, attacking accuracy, and combat efficiency, which were based on 77 sessions because of one missing match record. b represents the estimated difference in outcome (second cluster minus first cluster) derived from linear mixed-effects models adjusted for repeated measures within participants. CI: confidence interval.

**Supplementary Table 6.** Regression coefficients, 95% confidence intervals, and P-values from linear mixed-effects models comparing clusters on the NASA-TLX.

| Outcome | Comparison | Adjusted p-value | b (95 % CI) |
| --- | --- | --- | --- |
| NASA-TLX Total score (Figure 2D) | Cluster 1 vs Cluster 2 | 0.473 | 6.63 (-4.61, 17.88) |
|  | Cluster 1 vs Cluster 3 | 0.005 | 15.79 (3.74, 27.84) |
|  | Cluster 2 vs Cluster 3 | 0.115 | 9.15 (-1.41, 19.71) |
| TLX-subscales (Supplementary Figure 6D) |  |  |  |
| Mental demand | Cluster 1 vs Cluster 2 | 0.189 | 13.13 (-3.77, 30.02) |
|  | Cluster 1 vs Cluster 3 | < 0.001 | 30.29 (12.36, 48.22) |
|  | Cluster 2 vs Cluster 3 | 0.022 | 17.16 (1.84, 32.48) |
| Physical demand | Cluster 1 vs Cluster 2 | 0.456 | -7.64 (-20.41, 5.13) |
|  | Cluster 1 vs Cluster 3 | > 0.999 | -3.57 (-17.04, 9.90) |
|  | Cluster 2 vs Cluster 3 | > 0.999 | 4.07 (-7.22, 15.38) |
| Temporal demand | Cluster 1 vs Cluster 2 | > 0.999 | 0.01(-17.02, 17.04) |
|  | Cluster 1 vs Cluster 3 | 0.092 | 16.82 (-1.80, 35.44) |
|  | Cluster 2 vs Cluster 3 | 0.056 | 16.81 (-0.31, 33.93) |
| Own performance | Cluster 1 vs Cluster 2 | 0.089 | 13.78 (-1.37, 28.94) |
|  | Cluster 1 vs Cluster 3 | 0.137 | 13.91 (-2.74, 30.56) |
|  | Cluster 2 vs Cluster 3 | > 0.999 | 0.12 (-15.36, 15.61) |
| Effort | Cluster 1 vs Cluster 2 | > 0.999 | 6.23 (-10.15, 22.61) |
|  | Cluster 1 vs Cluster 3 | 0.274 | 12.22 (-5.10, 29.54) |
|  | Cluster 2 vs Cluster 3 | 0.983 | 5.99 (-8.66, 20.65) |
| Frustration | Cluster 1 vs Cluster 2 | 0.436 | 10.36 (-6.66, 27.38) |
|  | Cluster 1 vs Cluster 3 | 0.217 | 13.64 (-4.53, 31.80) |
|  | Cluster 2 vs Cluster 3 | > 0.999 | 3.28 (-12.48, 19.03) |

Analyses included 78 active play sessions from 13 participants. b represents the estimated difference in outcome (second cluster minus first cluster) derived from linear mixed-effects models adjusted for repeated measures within participants. CI, confidence interval.

**Supplementary Table 7.** Regression coefficients, 95% confidence intervals, and P-values from linear mixed-effects models comparing clusters on the Sports Flow Scale (Supplemental Figure 7A-B).

| Outcome | Comparison | Adjusted p-value | b (95 % CI) |
| --- | --- | --- | --- |
| Sports Flow Scale (SFS) total score | Cluster 1 vs Cluster 2 | >0.999 | 1.33 (-4.47, 7.12) |
|  | Cluster 1 vs Cluster 3 | 0.135 | 5.21 (-1.01, 11.45) |
|  | Cluster 2 vs Cluster 3 | 0.273 | 3.89 (-1.62, 9.40) |
| SFS-subscales |  |  |  |
| Fusion of action and perception | Cluster 1 vs Cluster 2 | >0.999 | 0.25 (-1.09, 1.59) |
|  | Cluster 1 vs Cluster 3 | >0.999 | 0.25 (-1.18, 1.68) |
|  | Cluster 2 vs Cluster 3 | >0.999 | 0.00 (-1.23, 1.24) |
| Focus on current task | Cluster 1 vs Cluster 2 | >0.999 | 0.31 (-0.66, 1.28) |
|  | Cluster 1 vs Cluster 3 | 0.004 | 1.42 (0.36, 2.48) |
|  | Cluster 2 vs Cluster 3 | 0.016 | 1.11(0.16, 2.06) |
| Sense of control | Cluster 1 vs Cluster 2 | >0.999 | -0.32 (-1.84, 1.20) |
|  | Cluster 1 vs Cluster 3 | >0.999 | 0.06 (-1.55, 1.68) |
|  | Cluster 2 vs Cluster 3 | >0.999 | 0.38 (-1.00, 1.76) |
| Loss of self-consciousness | Cluster 1 vs Cluster 2 | 0.017 | 1.77 (0.24, 3.30) |
|  | Cluster 1 vs Cluster 3 | 0.006 | 2.10 (0.48, 3.72) |
|  | Cluster 2 vs Cluster 3 | >0.999 | 0.32 (-1.05, 1.70) |
| Transformation of sense of time | Cluster 1 vs Cluster 2 | >0.999 | 0.35 (-1.66, 2.35) |
|  | Cluster 1 vs Cluster 3 | 0.027 | 2.39 (0.20, 4.58) |
|  | Cluster 2 vs Cluster 3 | 0.045 | 2.04 (0.03, 4.05) |
| Level of enjoyment in the activity | Cluster 1 vs Cluster 2 | >0.999 | 0.16 (-0.87, 1.19) |
|  | Cluster 1 vs Cluster 3 | 0.629 | 0.58 (-0.53, 1.69) |
|  | Cluster 2 vs Cluster 3 | 0.939 | 0.42 (-0.58, 1.41) |

Analyses included 78 active play sessions from 13 participants. b represents the estimated difference in outcome (second cluster minus first cluster) derived from linear mixed-effects models adjusted for repeated measures within participants. CI, confidence interval.

**Supplementary Table 8.**
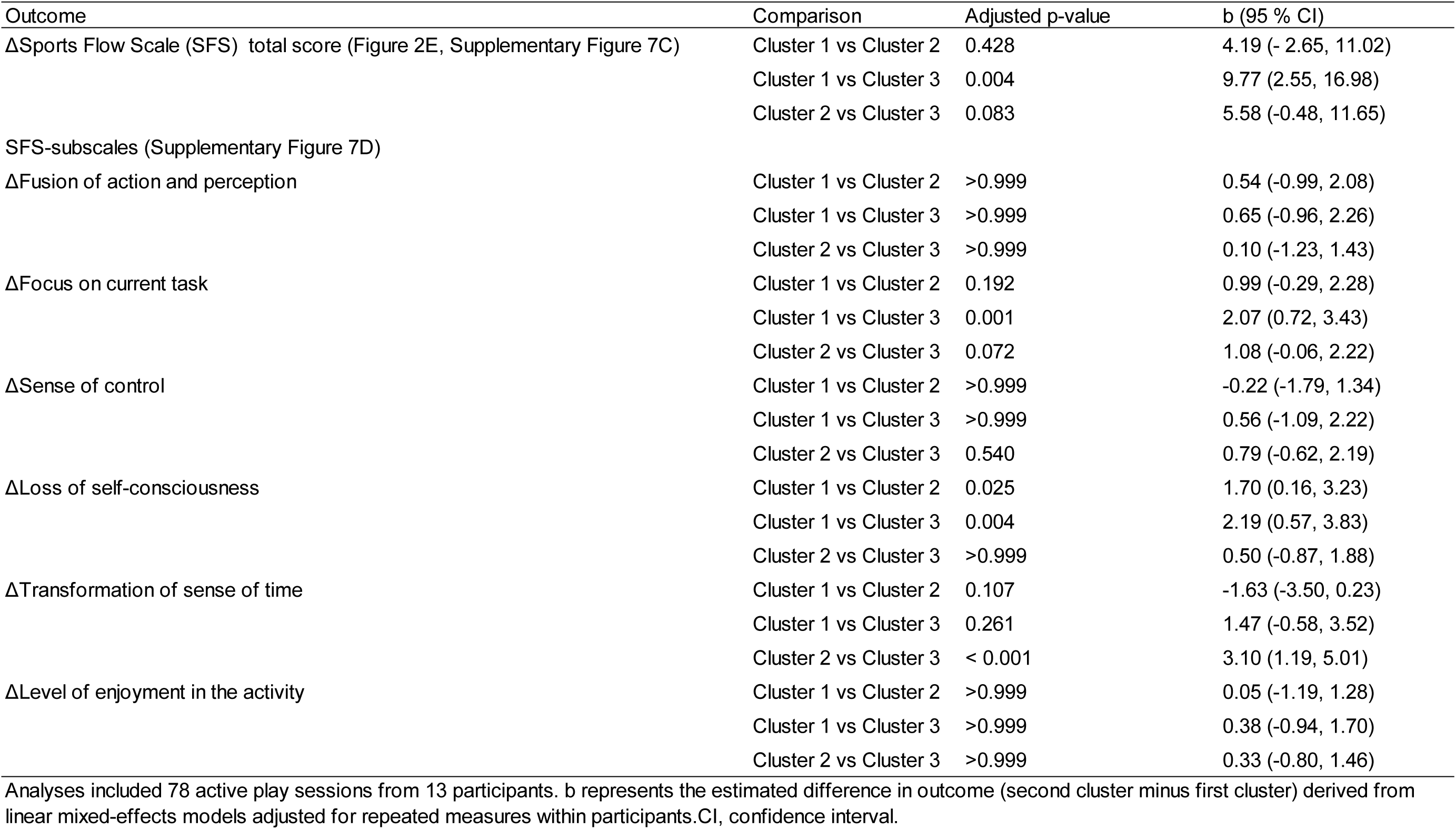
Regression coefficients, 95% confidence intervals, and P-values from linear mixed-effects models comparing clusters on changes in the Sports Flow Scale relative to the Watch condition.

| Outcome | Comparison | Adjusted p-value | b (95 % CI) |
| --- | --- | --- | --- |
| ΔSports Flow Scale (SFS) total score (Figure 2E, Supplementary Figure 7C) | Cluster 1 vs Cluster 2 | 0.428 | 4.19 (- 2.65, 11.02) |
|  | Cluster 1 vs Cluster 3 | 0.004 | 9.77 (2.55, 16.98) |
|  | Cluster 2 vs Cluster 3 | 0.083 | 5.58 (-0.48, 11.65) |
| SFS-subscales (Supplementary Figure 7D) |  |  |  |
| ΔFusion of action and perception | Cluster 1 vs Cluster 2 | >0.999 | 0.54 (-0.99, 2.08) |
|  | Cluster 1 vs Cluster 3 | >0.999 | 0.65 (-0.96, 2.26) |
|  | Cluster 2 vs Cluster 3 | >0.999 | 0.10 (-1.23, 1.43) |
| ΔFocus on current task | Cluster 1 vs Cluster 2 | 0.192 | 0.99 (-0.29, 2.28) |
|  | Cluster 1 vs Cluster 3 | 0.001 | 2.07 (0.72, 3.43) |
|  | Cluster 2 vs Cluster 3 | 0.072 | 1.08 (-0.06, 2.22) |
| ΔSense of control | Cluster 1 vs Cluster 2 | >0.999 | -0.22 (-1.79, 1.34) |
|  | Cluster 1 vs Cluster 3 | >0.999 | 0.56 (-1.09, 2.22) |
|  | Cluster 2 vs Cluster 3 | 0.540 | 0.79 (-0.62, 2.19) |
| ΔLoss of self-consciousness | Cluster 1 vs Cluster 2 | 0.025 | 1.70 (0.16, 3.23) |
|  | Cluster 1 vs Cluster 3 | 0.004 | 2.19 (0.57, 3.83) |
|  | Cluster 2 vs Cluster 3 | >0.999 | 0.50 (-0.87, 1.88) |
| ΔTransformation of sense of time | Cluster 1 vs Cluster 2 | 0.107 | -1.63 (-3.50, 0.23) |
|  | Cluster 1 vs Cluster 3 | 0.261 | 1.47 (-0.58, 3.52) |
|  | Cluster 2 vs Cluster 3 | < 0.001 | 3.10 (1.19, 5.01) |
| ΔLevel of enjoyment in the activity | Cluster 1 vs Cluster 2 | >0.999 | 0.05 (-1.19, 1.28) |
|  | Cluster 1 vs Cluster 3 | >0.999 | 0.38 (-0.94, 1.70) |
|  | Cluster 2 vs Cluster 3 | >0.999 | 0.33 (-0.80, 1.46) |

Analyses included 78 active play sessions from 13 participants. b represents the estimated difference in outcome (second cluster minus first cluster) derived from linear mixed-effects models adjusted for repeated measures within participants. CI, confidence interval.

**Supplementary Table 9.** Regression coefficients, 95% confidence intervals, and P-values from linear mixed-effects models comparing clusters on the cross-correlation analysis of pupil-heart dynamics (Supplementary Figure 8)

| Outcome | Comparison | Adjusted p-value | b (95 % CI) |
| --- | --- | --- | --- |
| Cross-correlation function | Cluster 1 vs Cluster 2 | 0.377 | 0.06 (-0.03, 0.17) |
|  | Cluster 1 vs Cluster 3 | >0.999 | -0.01 (-0.12, 0.10) |
|  | Cluster 2 vs Cluster 3 | 0.278 | -0.07 (-0.17, 0.03) |
| Timelag | Cluster 1 vs Cluster 2 | >0.999 | -0.40 (-1.64, 0.85) |
|  | Cluster 1 vs Cluster 3 | >0.999 | 0.45 (-0.90, 1.81) |
|  | Cluster 2 vs Cluster 3 | 0.313 | 0.85 (-0.40, 2.10) |

Analyses included 78 active play sessions from 13 participants. b represents the estimated difference in outcome (second cluster minus first cluster) derived from linear mixed-effects models adjusted for repeated measures within participants. CI, confidence interval.

**Supplementary Table 10.** Regression coefficients, 95% confidence intervals, and P-values from linear mixed-effects models comparing clusters on the cross reccurence quantification analysis of pupil-heart dynamics (Figure 3B-E, Supplementary Figure 9)

| Outcome | Comparison | Adjusted p-value | b (95 % CI) |
| --- | --- | --- | --- |
| Radius | Cluster 1 vs Cluster 2 | 0.033 | 0.04 (0.00, 0.09) |
|  | Cluster 1 vs Cluster 3 | 0.020 | 0.05 (0.01, 0.10) |
|  | Cluster 2 vs Cluster 3 | >0.999 | 0.01 (-0.03, 0.05) |
| Determinism | Cluster 1 vs Cluster 2 | >0.999 | 0.01 (-0.06, 0.07) |
|  | Cluster 1 vs Cluster 3 | >0.999 | -0.01 (-0.08, 0.06) |
|  | Cluster 2 vs Cluster 3 | >0.999 | -0.02 (-0.08, 0.04) |
| Average diagonal line length | Cluster 1 vs Cluster 2 | >0.999 | -0.02 (-0.17, 0.13) |
|  | Cluster 1 vs Cluster 3 | >0.999 | -0.05 (-0.22, 0.11) |
|  | Cluster 2 vs Cluster 3 | >0.999 | -0.03 (-0.18, 0.12) |
| Maximal diagonal line length | Cluster 1 vs Cluster 2 | 0.396 | -0.74 (-1.91, 0.43) |
|  | Cluster 1 vs Cluster 3 | 0.004 | -1.74 (-3.04, -0.45) |
|  | Cluster 2 vs Cluster 3 | 0.149 | -1.01 (-2.24, 0.22) |
| Diagonal entropy | Cluster 1 vs Cluster 2 | >0.999 | -0.04 (-0.15, 0.08) |
|  | Cluster 1 vs Cluster 3 | 0.460 | -0.07 (-0.20, 0.05) |
|  | Cluster 2 vs Cluster 3 | >0.999 | -0.04 (-0.16, 0.08) |
| Laminarity | Cluster 1 vs Cluster 2 | >0.999 | -0.01 (-0.08, 0.06) |
|  | Cluster 1 vs Cluster 3 | >0.999 | -0.02 (-0.10, 0.05) |
|  | Cluster 2 vs Cluster 3 | >0.999 | -0.02 (-0.08, 0.05) |
| Average vertical line length | Cluster 1 vs Cluster 2 | >0.999 | 0.26 (-0.55, 1.06) |
|  | Cluster 1 vs Cluster 3 | 0.417 | 0.54 (-0.33, 1.40) |
|  | Cluster 2 vs Cluster 3 | >0.999 | 0.28 (-0.50, 1.06) |
| Maximal vertical line length | Cluster 1 vs Cluster 2 | >0.999 | 2.75 (-5.72, 11.21) |
|  | Cluster 1 vs Cluster 3 | >0.999 | -2.43 (-11.69, 6.83) |
|  | Cluster 2 vs Cluster 3 | 0.438 | -5.17 (-13.70, 3.35) |
| Vertical entropy | Cluster 1 vs Cluster 2 | >0.999 | 0.05 (-0.20, 0.30) |
|  | Cluster 1 vs Cluster 3 | 0.530 | 0.15 (-0.12, 0.42) |
|  | Cluster 2 vs Cluster 3 | 0.897 | 0.10 (-0.13, 0.34) |

Analyses included 78 active play sessions from 13 participants. b represents the estimated difference in outcome (second cluster minus first cluster) derived from linear mixed-effects models adjusted for repeated measures within participants. CI, confidence interval.

**Supplementary Table 11.** Direct and indirect effects of pupil–heart synchronization on in-game performance outcomes mediated by subjective workload in the pooled session analysis (NASA-TLX) (Figure 4; Supplementary Figure 10)

| Outcome | Direct effect |  | Indirect effect |  |
| --- | --- | --- | --- | --- |
|  | b (95%CI) | p-value | b (95%CI) | p-value |
| Number of knockouts | 3.33 (-2.01, 8.67) | 0.221 | -2.59 (-5.00, -0.17) | 0.036 |
| Combat efficiency | -1.81 (-6.43, 2.82) | 0.444 | -2.69 (-5.13, -0.25) | 0.030 |
| Damage dealt | 119.94 (-282.29, 522.17) | 0.559 | -324.12 (-612.47, -35.76) | 0.028 |
| Damage taken | 556.65 (65.19, 1048.10) | 0.026 | 193.47 (24.19, 362.75) | 0.025 |
| Attacking accuracy | 18.04 (-7.62, 43.71) | 0.168 | -9.27 (-20.41, 1.86) | 0.103 |

Analyses included 78 active play sessions from 13 participants, except for damage taken, attacking accuracy, and combat efficiency, which were based on 77 sessions because of one missing match record. Pupil–heart synchronization was indexed by CRQA radius (lower values indicate stronger synchronization). Indirect effects were estimated through NASA-TLX total score using mediation analyses.

## References

1. D. W. Franklin, D. M. Wolpert, Computational mechanisms of sensorimotor control. Neuron 72, 425–442 (2011).

2. J. P. Gallivan, C. S. Chapman, D. M. Wolpert, J. R. Flanagan, Decision-making in sensorimotor control. Nat Rev Neurosci 19, 519–534 (2018).

3. B. M. Herbert, P. Ulbrich, R. Schandry, Interoceptive sensitivity and physical effort: implications for the self-control of physical load in everyday life. Psychophysiology 44, 194–202 (2007).

4. B. Mouatt et al., What I see and what I feel: the influence of deceptive visual cues and interoceptive accuracy on affective valence and sense of effort during virtual reality cycling. PeerJ 11, e16095 (2023).

5. E. Kokkinara, K. Kilteni, K. J. Blom, M. Slater, First Person Perspective of Seated Participants Over a Walking Virtual Body Leads to Illusory Agency Over the Walking. Sci Rep 6, 28879 (2016).

6. G. Kong, K. He, K. Wei, Sensorimotor experience in virtual reality enhances sense of agency associated with an avatar. Conscious Cogn 52, 115–124 (2017).

7. T. Matsui et al., Cognitive decline with pupil constriction independent of subjective fatigue during prolonged esports across player expertise levels. Computers in Human Behavior 156 (2024).

8. M. Kellmann et al., Recovery and Performance in Sport: Consensus Statement. Int J Sports Physiol Perform 13, 240–245 (2018).

9. C. Maslach, W. B. Schaufeli, M. P. Leiter, Job Burnout. Annual Review of Psychology 52, 397–422 (2001).

10. B. Bediou et al., Meta-analysis of action video game impact on perceptual, attentional, and cognitive skills. Psychol Bull 144, 77–110 (2018).

11. K. Cahill, M. Dhamala, Structurally constrained functional connectivity reveals efficient visuomotor decision-making mechanisms in action video gamers. Sci Rep 15, 43301 (2025).

12. I. Pedraza-Ramirez, L. Musculus, M. Raab, B. Ramaker, S. Laborde, Zooming in on decision making in esports: Exploring the perceptions of expert players and coaches. *Sport*, Exercise, and Performance Psychology 14, 335–351 (2025).

13. M. J. Koepp et al., Evidence for striatal dopamine release during a video game. Nature 393, 266–268 (1998).

14. J. Nakamura, M. Csikszentmihalyi, "The Concept of Flow" in Handbook of Positive Psychology, C. R. Snyder, S. J. Lopez, Eds. (Oxford University Press, New York, NY, 2002), pp. 89-105.

15. L. Harmat et al., Physiological correlates of the flow experience during computer game playing. Int J Psychophysiol 97, 1–7 (2015).

16. D. J. Harris, K. L. Allen, S. J. Vine, M. R. Wilson, A systematic review and meta-analysis of the relationship between flow states and performance. International Review of Sport and Exercise Psychology 16, 693–721 (2021).

17. J. Palomäki et al., The link between flow and performance is moderated by task experience. Computers in Human Behavior 124 (2021).

18. H. Miao, H. He, X. Hou, J. Wang, L. Chi, Cognitive expertise in esport experts: a three-level model meta-analysis. PeerJ 12, e17857 (2024).

19. M. A. Pluss et al., Perceptual-motor Abilities Underlying Expertise in Esports Journal of Expertise 3, 133–143 (2020).

20. H. D. Critchley, N. A. Harrison, Visceral influences on brain and behavior. Neuron 77, 624–638 (2013).

21. G. Forte, F. Favieri, M. Casagrande, Heart Rate Variability and Cognitive Function: A Systematic Review. Front Neurosci 13, 710 (2019).

22. J. F. Thayer, A. L. Hansen, E. Saus-Rose, B. H. Johnsen, Heart rate variability, prefrontal neural function, and cognitive performance: the neurovisceral integration perspective on self-regulation, adaptation, and health. Ann Behav Med 37, 141–153 (2009).

23. A. Bashan, R. P. Bartsch, J. W. Kantelhardt, S. Havlin, P. Ivanov, Network physiology reveals relations between network topology and physiological function. Nat Commun 3, 702 (2012).

24. A. W. Corcoran, A. Le Coz, J. Hohwy, T. Andrillon, When your heart isn’t in it anymore: cardiac correlates of task disengagement. Commun Biol 8, 1646 (2025).

25. D. Candia-Rivera, M. Vidailhet, M. Chavez, F. De Vico Fallani, A framework for quantifying the coupling between brain connectivity and heartbeat dynamics: Insights into the disrupted network physiology in Parkinson’s disease. Hum Brain Mapp 45, e26668 (2024).

26. T. W. P. Janssen et al., Opportunities and Limitations of Mobile Neuroimaging Technologies in Educational Neuroscience. Mind Brain Educ 15, 354–370 (2021).

27. S. Joshi, Y. Li, R. M. Kalwani, J. I. Gold, Relationships between Pupil Diameter and Neuronal Activity in the Locus Coeruleus, Colliculi, and Cingulate Cortex. Neuron 89, 221–234 (2016).

28. P. R. Murphy, R. G. O’Connell, M. O’Sullivan, I. H. Robertson, J. H. Balsters, Pupil diameter covaries with BOLD activity in human locus coeruleus. Human Brain Mapping 35, 4140–4154 (2014).

29. M. Behnke, A. Hase, L. D. Kaczmarek, P. Freeman, Blunted cardiovascular reactivity may serve as an index of psychological task disengagement in the motivated performance situations. Sci Rep 11, 18083 (2021).

30. O. Durcan, P. Holland, J. Bhattacharya, A framework for neurophysiological experiments on flow states. Commun Psychol 2, 66 (2024).

31. Y. Liu, S. Narasimhan, B. J. Schriver, Q. Wang, Perceptual Behavior Depends Differently on Pupil-Linked Arousal and Heartbeat Dynamics-Linked Arousal in Rats Performing Tactile Discrimination Tasks. Front Syst Neurosci 14, 614248 (2021).

32. K. Watanabe, N. Saijo, S. Minami, M. Kashino, The effects of competitive and interactive play on physiological state in professional esports players. Heliyon 7, e06844 (2021).

33. D. Weibel, B. Wissmath, S. Habegger, Y. Steiner, R. Groner, Playing online games against computer- vs. human-controlled opponents: Effects on presence, flow, and enjoyment. Computers in Human Behavior 24, 2274–2291 (2008).

34. L. Nacke, C. A. Lindley, Flow and Immersion in First-Person Shooters: Measuring the player’s gameplay experience. Conference: Proceedings of the 2008 Conference on Future Play, 81-88.

35. M. Shehata et al., Team Flow Is a Unique Brain State Associated with Enhanced Information Integration and Interbrain Synchrony. eNeuro 8 (2021).

36. L. Li, D. M. Smith, Neural Efficiency in Athletes: A Systematic Review. Front Behav Neurosci 15, 698555 (2021).

37. A. C. Neubauer, A. Fink, Intelligence and neural efficiency. Neurosci Biobehav Rev 33, 1004–1023 (2009).

38. V. Bianco, F. Di Russo, R. L. Perri, M. Berchicci, Different proactive and reactive action control in fencers’ and boxers’ brain. Neuroscience 343, 260–268 (2017).

39. T. S. Braver, The variable nature of cognitive control: a dual mechanisms framework. Trends Cogn Sci 16, 106–113 (2012).

40. C. Swann et al., Psychological States Underlying Excellent Performance in Sport: Toward an Integrated Model of Flow and Clutch States. Journal of Applied Sport Psychology 29, 375–401 (2017).

41. G. Aston-Jones, J. D. Cohen, An integrative theory of locus coeruleus-norepinephrine function: Adaptive gain and optimal performance. Annual Review of Neuroscience 28, 403–450 (2005).

42. S. N. Meissner et al., Self-regulating arousal via pupil-based biofeedback. Nat Hum Behav 8, 43–62 (2024).

43. H. Lu, D. van der Linden, A. B. Bakker, Changes in pupil dilation and P300 amplitude indicate the possible involvement of the locus coeruleus-norepinephrine (LC-NE) system in psychological flow. Sci Rep 13, 1908 (2023).

44. A. Fourcade et al., Linking brain-heart interactions to emotional arousal in immersive virtual reality. Psychophysiology 61, e14696 (2024).

45. Y. Ebara, T. Sakai, Organization of terms related to mental methods in early modern swordsmanship with a focus on the early Shinkage-Yagyu school. Research Journal of Physical Arts 2 (1995).

46. M. Takenaka, Y. Aida, S. Y., Outstanding female kendo players’ practical knowledge focusing on tactics of the most accomplished. Japan Journal of Physical Education 69, 139–149 (2024).

47. R. Mazziotti et al., MEYE: Web App for Translational and Real-Time Pupillometry. eNeuro 8 (2021).

48. M. Hassan et al., Augmenting the Sense of Social Presence in Online Video Games Through the Sharing of Biosignals. IEEE Access 12, 98977–98989 (2024).

49. S. Cotterill, Pre-performance routines in sport: current understanding and future directions. International Review of Sport and Exercise Psychology 3, 132–153 (2010).

50. M. J. Duncan et al., Effects of increasing and decreasing physiological arousal on anticipation timing performance during competition and practice. Eur J Sport Sci 16, 27–35 (2016).

51. R. Kuwamizu et al., Pupil-linked arousal with very light exercise: pattern of pupil dilation during graded exercise. J Physiol Sci 72, 23 (2022).

52. S. Takahashi, W. Kosugi, S. Mizuno, T. Matsui, Sparkling water consumption mitigates cognitive fatigue during prolonged esports play. Computers in Human Behavior Reports 21 (2026).

53. H. van Steenbergen, T. F. Wilderjans, G. P. H. Band, S. T. Nieuwenhuis, Boosting arousal and cognitive performance through alternating posture: Insights from a multi-method laboratory study. Psychophysiology 61, e14634 (2024).

## Appendix: References

1. T. Matsui et al., Cognitive decline with pupil constriction independent of subjective fatigue during prolonged esports across player expertise levels. Computers in Human Behavior 156 (2024).

2. T. Monma et al., Prevalence and Associated Factors of Physical Complaints Among Japanese Esports Players: A Cross-Sectional Study. Cureus 16, e66496 (2024).

3. N. Murase, Validity and reliability of Japanese version of International Physical Activity Questionnaire. Journal of Health and Welfare Statistics 49, 1–9 (2002).

4. T. Yagi, T. Sakairi, Subjective Arousal and Experience of Flow during Positive Sports Event. The Japanese Journal of Health Psychology 22, 24–32 (2009).

5. S. G. Hart, L. E. Staveland, "Development of NASA-TLX (Task Load Index): Results of Empirical and Theoretical Research" in Human Mental Workload. (1988), 10.1016/s0166-4115(08)62386-9, pp. 139-183.

6. S. Miyake, M. Kumashiro, Subjective mental workload assessment technique-an introduction to NASA-TLX and SWAT and a proposal of simple scoring methods. The Japanese Journal of Ergonomics 29, 399–408 (1993).

7. J. MacQueen (1967) Some methods for classification and analysis of multivariate observations. in Proceedings of the Fifth Berkeley Symposium on Mathematical Statistics and Probability (University of California Press, Berkeley, CA), pp 281-297.

8. M. I. Coco, R. Dale, Cross-recurrence quantification analysis of categorical and continuous time series: an R package. Front Psychol 5, 510 (2014).

9. S. Wallot, G. Leonardi, Analyzing Multivariate Dynamics Using Cross-Recurrence Quantification Analysis (CRQA), Diagonal-Cross-Recurrence Profiles (DCRP), and Multidimensional Recurrence Quantification Analysis (MdRQA) - A Tutorial in R. Front Psychol 9, 2232 (2018).

10. X. Gu, X. Liang, S. Zhao, Agudamu, Y. Zhang, Team Synchrony Under Fire: Heart Rate Recurrence Reveals a Biosignature of Victory in Elite Esports. Med Sci Sports Exerc 58, 572–580 (2026).

11. A. M. Fraser, H. L. Swinney, Independent coordinates for strange attractors from mutual information. Phys Rev A Gen Phys 33, 1134–1140 (1986).

12. M. B. Kennel, R. Brown, H. D. Abarbanel, Determining embedding dimension for phase-space reconstruction using a geometrical construction. Phys Rev A 45, 3403–3411 (1992).

13. N. Marwan, M. Carmenromano, M. Thiel, J. Kurths, Recurrence plots for the analysis of complex systems. Physics Reports 438, 237–329 (2007).

14. K. H. Kraemer, R. V. Donner, J. Heitzig, N. Marwan, Recurrence threshold selection for obtaining robust recurrence characteristics in different embedding dimensions. Chaos 28, 085720 (2018).

15. T. Matsui et al., Endogenous oxytocin-linked heart-rate synchrony and social connection during live sports spectatorship. Translational Psychiatry 10.1038/s41398-026-04265-2 (2026).

